# The landscape of drug sensitivity and resistance in sarcoma

**DOI:** 10.1101/2023.05.25.542375

**Authors:** Ahmad Al Shihabi, Peyton J Tebon, Huyen Thi Lam Nguyen, Jomjit Chantharasamee, Sara Sartini, Ardalan Davarifar, Alexandra Y Jensen, Miranda Diaz-Infante, Hannah Cox, Alfredo Enrique Gonzalez, Summer Swearingen, Nasrin Tavanaie, Sarah Dry, Arun Singh, Bartosz Chmielowski, Joseph G. Crompton, Anusha Kalbasi, Fritz C Eilber, Francis Hornicek, Nicholas Bernthal, Scott D Nelson, Paul C Boutros, Noah Federman, Jane Yanagawa, Alice Soragni

## Abstract

Sarcomas are a family of rare malignancies composed of over 100 distinct histological subtypes. The rarity of sarcoma poses significant challenges in conducting clinical trials to identify effective therapies, to the point that many rarer subtypes of sarcoma do not have standard-of-care treatment. Even for established regimens, there can be substantial heterogeneity in responses. Overall, novel, personalized approaches for identifying effective treatments are needed to improve patient outcomes. Patient-derived tumor organoids (PDTOs) are clinically relevant models representative of the physiological behavior of tumors across an array of malignancies. Here, we use PDTOs as a tool to better understand the biology of individual tumors and characterize the landscape of drug resistance and sensitivity in sarcoma. We collected n=194 specimens from n=126 sarcoma patients, spanning 24 distinct subtypes. We characterized PDTOs established from over 120 biopsy, resection, and metastasectomy samples. We leveraged our organoid high-throughput drug screening pipeline to test the efficacy of chemotherapeutics, targeted agents, and combination therapies, with results available within a week from tissue collection. Sarcoma PDTOs showed patient-specific growth characteristics and subtype-specific histopathology. Organoid sensitivity correlated with diagnostic subtype, patient age at diagnosis, lesion type, prior treatment history, and disease trajectory for a subset of the compounds screened. We found 90 biological pathways that were implicated in response to treatment of bone and soft tissue sarcoma organoids. By comparing functional responses of organoids and genetic features of the tumors, we show how PDTO drug screening can provide an orthogonal set of information to facilitate optimal drug selection, avoid ineffective therapies, and mirror patient outcomes in sarcoma. In aggregate, we were able to identify at least one effective FDA-approved or NCCN-recommended regimen for 59% of the specimens tested, providing an estimate of the proportion of immediately actionable information identified through our pipeline.

**Highlights:**

- Standardized organoid culture preserve unique sarcoma histopathological features
- Drug screening on patient-derived sarcoma organoids provides sensitivity information that correlates with clinical features and yields actionable information for treatment guidance
- High-throughput screenings provide orthogonal information to genetic sequencing
- Sarcoma organoid response to treatment correlates with patient response to therapy
- Large scale, functional precision medicine programs for rare cancers are feasible within a single institution

## Introduction

Sarcomas are a family of rare and heterogeneous tumors of mesenchymal origin^1^. Clinically, these tumors primarily arise in bone and soft tissue and disproportionately impact young patients^1–3^. Despite a low incidence, with ∼13,000 soft tissue sarcoma and ∼4,000 bone sarcoma diagnoses annually in the United States^3^, fatalities remain high. Bone sarcomas, for instance, are the third leading cause of cancer deaths in patients under 20 years old^3^. The treatment regimen varies greatly by disease subtype and stage; therapeutic options include surgical resection, chemotherapy, targeted systemic therapy, and radiotherapy in certain cases^4,5^. Despite significant advances for specific subtypes, current treatment approaches are rarely curative and contribute to aggregate overall 5-year survival rates of ∼65% in soft tissue sarcoma and ∼50-60% for bone cancers^6^.

The heterogeneity of sarcoma manifests in over 100 distinct subtypes. Diversity is observed across and within sarcoma diagnosis: for instance, there are 9 distinct osteosarcoma subtypes reported^7^. About a third of sarcoma cases are driven by specific chromosomal fusions, such as subsets of Ewing sarcoma, synovial sarcoma, infantile fibrosarcoma and rhabdomyosarcoma^8^. Other key genetic events across bone and soft tissue sarcomas are thought to impact cell cycle regulation, growth factor signaling, and angiogenesis^9^. Specific high-prevalence mutations include RTK/RAS driver mutations among epithelioid sarcomas^10^, CDK aberrations in liposarcomas^10^, and PI3K mutations in PEComa and myxoid liposarcomas^10^. This vast heterogeneity compounds the challenge to identify effective regimens for this family of rare and ultra-rare cancers and contributes to persistently low survival rates.

Precision medicine approaches are attracting increasing interest as tools to identify actionable characteristics and improve outcomes on a per-patient basis^11^. Technologies such as next-generation sequencing (NGS) and immunohistochemistry are widely used to identify molecular alterations and druggable targets^11–14^. In the case of fusion-positive sarcoma, most of the aberrant oncogenes cannot be targeted directly, with the notable exception of NTRK^8^. A recent study on over 6’000 bone and soft tissue sarcomas found an average of ∼42% of tumors harboring actionable alterations by NGS^11^. Despite this, few sarcoma patients show clinical benefit when treated with drugs selected via genomic precision medicine, as demonstrated in many clinical trials over the past 10 years^9,11,15,16^.

Because of this genetic diversity and the limited efficacy of chemotherapeutic and targeted agents selected through both conventional or precision medicine, there is a dire need to identify alternative approaches to systematically evaluate the landscape of drug sensitivity and resistance in sarcoma and identify individualized therapeutic solutions. Here, we leverage patient-derived tumor organoid-based functional assays as an alternative yet complementary approach to genetic-based precision medicine in sarcoma^17^. Patient-derived organoids (PDTOs) are ideally suited to fill this gap, as they are tractable and predictive of response to treatment, at least for patients with various types of epithelial cancers^18–25^. The development of sarcoma PDTOs has lagged, with limited applications so far^26,27^. We have procured n=194 specimens from 126 patients undergoing biopsies or surgical resections at UCLA Health hospitals and successfully generated PDTOs from over 121 samples originating from primary, recurrent and metastatic bone and soft tissue sarcomas. Here we describe our pipeline for procuring and generating sarcoma PDTOs, which includes histopathology to characterize both the parent tumor and PDTOs as well as sequencing and high throughput drug testing leveraging our rapid 3D organoid screening platform^17,26,28–30^. The drug sensitivity and resistance patterns observed across 24 sarcoma types pinpointed both subtype-specific and patient-specific vulnerabilities.

## Results

### Clinical characteristics of sarcoma patients and samples

We collected a total of n=194 sarcoma specimens from 126 patients treated at UCLA for a sarcoma diagnosis between February 2018 and May 2022 (**Figure 1**). Tissue was obtained from biopsies (n=11) or surgical resections (n=183) of primary, recurrent, and metastatic lesions (**Figure 1A and 1B**). The patient population was majority adult (n=63/126) with 35% adolescent and young adults (AYA, n=45/126) and 13% pediatric patients (n=18/126) at time of diagnosis (**Figure 1B, Figure S1A and S1B**). Patients were 62% male (78/126) and 38% female (48/126), which closely resembles the proportion of incidence of bone and soft tissue sarcomas in the United States (57% male, 43% female)^31^. 15% identified as Asian, 5% as Black, 1% as Pacific Islander/Samoan and 18% as Other.

**Figure 1.**
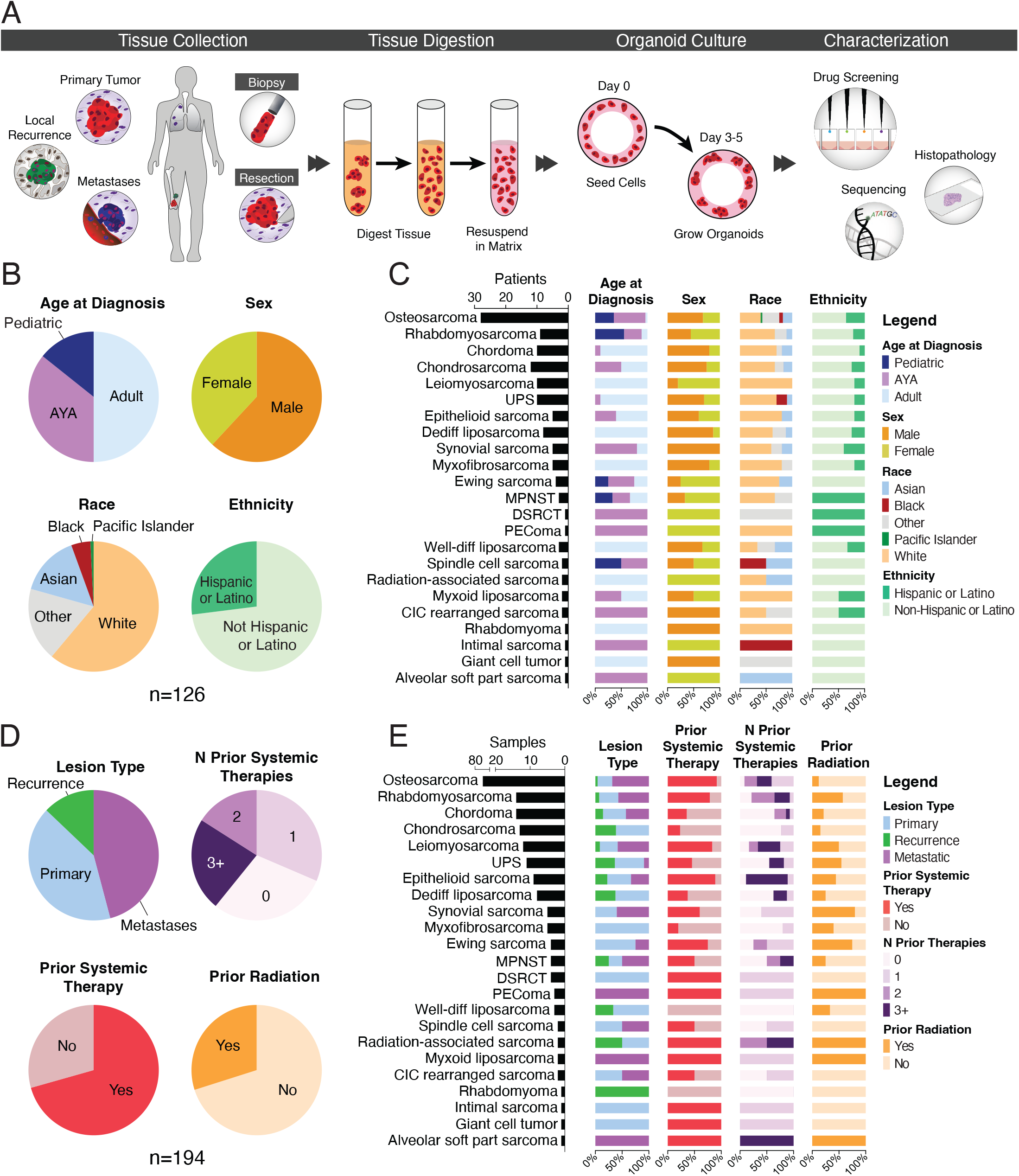
Overview of the patient-derived tumor organoid pipeline, patient demographics and sample characteristics. (A) Tissue is collected from biopsies or surgical resections of bone and soft tissue sarcomas. Organoids are generated by digesting the tissue and culturing the harvested cells in a 3D matrix. Organoids are molecularly and functionally profiled. (B) Demographics of pan-sarcoma study patients. Pediatric: 0-14 years old, AYA: 15-39 years old, adults: 40 years old and above. (C) Demographics divided by diagnosis. (D) Clinical characteristics of tumors from which tissue was collected. Prior systemic therapies include chemotherapy, targeted agents, and immunotherapy. (E) Tumor characteristics divided by disease subtype.

Our study includes tumor samples from 24 distinct subtypes of bone and soft tissue sarcomas (**Figure 1B and 1C**). The most common diagnosis in this cohort is osteosarcoma, for which we collected n=73 samples from 28 patients, followed by chordoma (n=14 samples from 10 patients) and chondrosarcoma (n=13 samples from 12 patients). For soft tissue sarcoma, the majority of cases were leiomyosarcoma (n=12 samples from 10 patients) and undifferentiated pleomorphic sarcoma (n=11 samples from 10 patients, **Figure 1C**). The proportion of patients enrolled reflects the proportional incidences of the major sarcoma subtypes, with the exceptions of chordoma (overrepresented in our study) and leiomyosarcoma (underrepresented in our study)^32^.

The majority of samples procured were from metastases (46%), followed by primary tumors (41%), and recurrent tumors (13%). In 21% of cases (27 patients), we collected multiple specimens from the same patient, either from different anatomical locations (12/27), longitudinally (21/27) or both (6/27). Regardless of tumor type, most samples (71%) were exposed to various systemic regimens, with 23% (n=45/194) treated with ≥ 3 prior lines of systemic therapy while 30% had a history of radiation exposure (**Figure 1D**). The median interval between last systemic treatment and sample procurement was one month for the 137 pre-treated specimens (range: 1-480 mo, **Figure S1D**), and 12 months for the 59 irradiated samples (range: 1-384 mo, **Figure S1C**). The proportion of tumors treated with each modality varied across sarcoma subtypes, reflecting the heterogeneity in the clinical management of each unique diagnosis (**Figure 1E**)^4,5^. For instance, myxoid liposarcomas are commonly known for their radiosensitivity and treated with radiation pre-operatively as standard of care, which is reflected in our cohort (**Figure 1E**)^33^.

### Establishment of soft tissue and bone sarcoma PDTOs

The specimens included in this study were procured through the UCLA Pathology Department and expedited to the lab for processing following our published protocols as described in the Methods section (**Figure 1A**)^17,26,28–30^. Given that there is no a priori sorting of tumor cells, we performed a detailed histopathological analysis of all organoids and parental tissue to verify that tumor cells were present and all main features were maintained ex vivo^26,29^ (**Figure 1A and 2**). We observed diverse histopathological features across the sarcoma subtypes, such as small round blue cells in Ewing sarcoma, vacuolated cells with round nuclei in chordoma, and spindle cells in rhabdomyosarcoma, some types of osteosarcomas and malignant peripheral nerve sheath tumors (MPNSTs, **Figure 2**). Across all subtypes, the organoids largely recapitulated all the salient features of the native tissue (**Figure 2**). For instance, both organoids generated from small cell osteosarcoma and Ewing sarcoma showed the characteristic small, round nuclei while heterogeneous nucleus sizes and structures were found in undifferentiated pleomorphic sarcomas (UPS). Consistent with our prior work^26^, chordoma organoids grew in clusters with extensive production of vacuoles as seen in the parental tumors (**Figure 2**).

**Figure 2.**
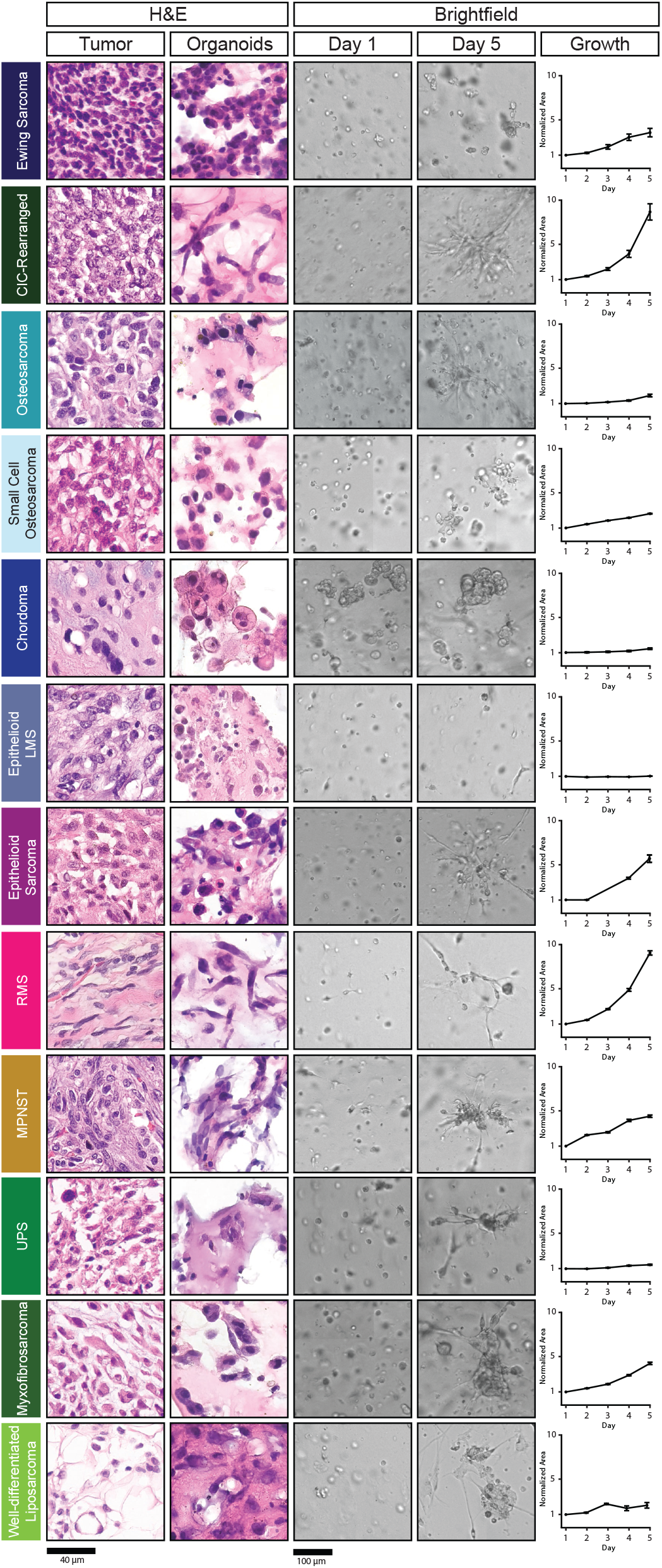
Sarcoma organoids grow in culture and recapitulate key morphological features of the parental tumors. Representative images of sarcomas and corresponding organoids stained with H&E (columns 1 and 2). Representative brightfield images of the same sarcoma cells in culture on Day 1 (column 3) and Day 5 (column 4). Growth was tracked over time by segmenting in-focus organoids in the brightfield images using a machine learning-based pipeline and by normalizing the cross-sectional area covered by organoids to that measured on the first day of culture. Scale bars: 40 μM for H&E images, 100 μM for brightfield pictures.

In addition to histopathology, we employed serial brightfield imaging to quantitatively assess the growth of organoids ex vivo^26^ (**Figure 2**). Plates were imaged once every 24 hours and growth rates quantified by applying a U-net machine learning-based segmentation algorithm that quantifies the cross-sectional area occupied by organoids^26,34^ (**Figure 2**). Organoids derived from UPS, LMS, and chordoma demonstrated minimal increases in normalized area over five days. This aligns with our previous findings in chordoma organoids, where limited proliferation and cell rearrangement were observed in PDTOs from multiple patients^26^. In contrast, PDTOs from patients diagnosed with myxofibrosarcoma, RMS, epithelioid sarcoma, and CIC-rearranged sarcoma exhibited exponential growth, characterized by cell aggregation and proliferation, leading to extensive networks of multicellular clusters with large cross-sectional areas (**Figure 2**). Ewing sarcoma organoids displayed an initial phase of exponential growth followed by a plateau, yet maintained their round morphology—a unique characteristic observed in these organoids. Lastly, a subset of PDTOs demonstrated near-linear growth dynamics (**Figure 2**).

Overall, our data demonstrates that PDTOs established from diverse soft tissue and bone sarcoma subtypes maintain the key histological characteristics of the tumor of origin while exhibiting diverse growth dynamics ex vivo. This suggests that PDTOs maintain sarcoma subtype-specific and patient-specific phenotypes within our standardized approach for organoid culture.

### Sarcoma PDTO screening pipeline

We procured n=194 samples total that were expedited to the lab and processed for downstream characterization and screening (**Figure S2A**). Multiple samples were collected from 27 patients. Of all collected samples, 21 had insufficient cell number for further analyses, with less than 250’000 viable cells recovered. Four additional samples had low cell numbers but were seeded for expansion and passaging yet failed to grow. Lastly, two samples were excluded due to contamination by ink marking during the pathology workflow and three were used for different studies. A total of 40 samples were frozen, including 19 that were procured during COVID-related shutdowns and interruptions. The remaining n=124 samples corresponding to 80% of all samples that were processed, were used to generate mini-rings^17,26,28–30^ in 96 well plates for high throughput drug screening. These encompassed 21 different sarcoma diagnoses. Ten samples showed no growth as measured by low luminescence values post-ATP-release assay, for an overall organoid take rate of ∼93% considering all samples we attempted to screen or grow and expand (**Figure S2A**).

We used Z’ factor and robust Z’ factor parameters to assess the quality of drug screening results^35,36^. To reduce the likelihood of false positives and false negatives, we included plates with a Z’ factor or robust Z’ factor greater than 0.2 in downstream analyses (**Figure S2B and S2C**). Of the n=114 samples that were screened and exhibited robust growth, 16 did not meet our quality control cutoffs (**Figure S2**).

The drug library utilized in our study includes over 400 compounds and drug combinations in different stages of clinical and pre-clinical development (**Figure 3A-B**)^26,28,29^. For each sample, we customized a panel of drugs to be screened based on sample characteristics, histology, a priori patient-specific information, and clinical data. Our selection prioritized FDA-approved and NCCN-recommended regimes specific to each sarcoma subtype or other cancers and included molecules targeting suspected or confirmed genetic alterations for each case. We also incorporated the anticipated treatment plan and drug combinations currently being evaluated in clinical trials. On average, we tested n=117 drugs/sample (range: 6-423), depending on the number of viable cells extracted and drug panel of interest. This approach allowed for a comprehensive evaluation of potential therapeutic options, tailored to the individual characteristics and clinical context of each sample.

**Figure 3.**
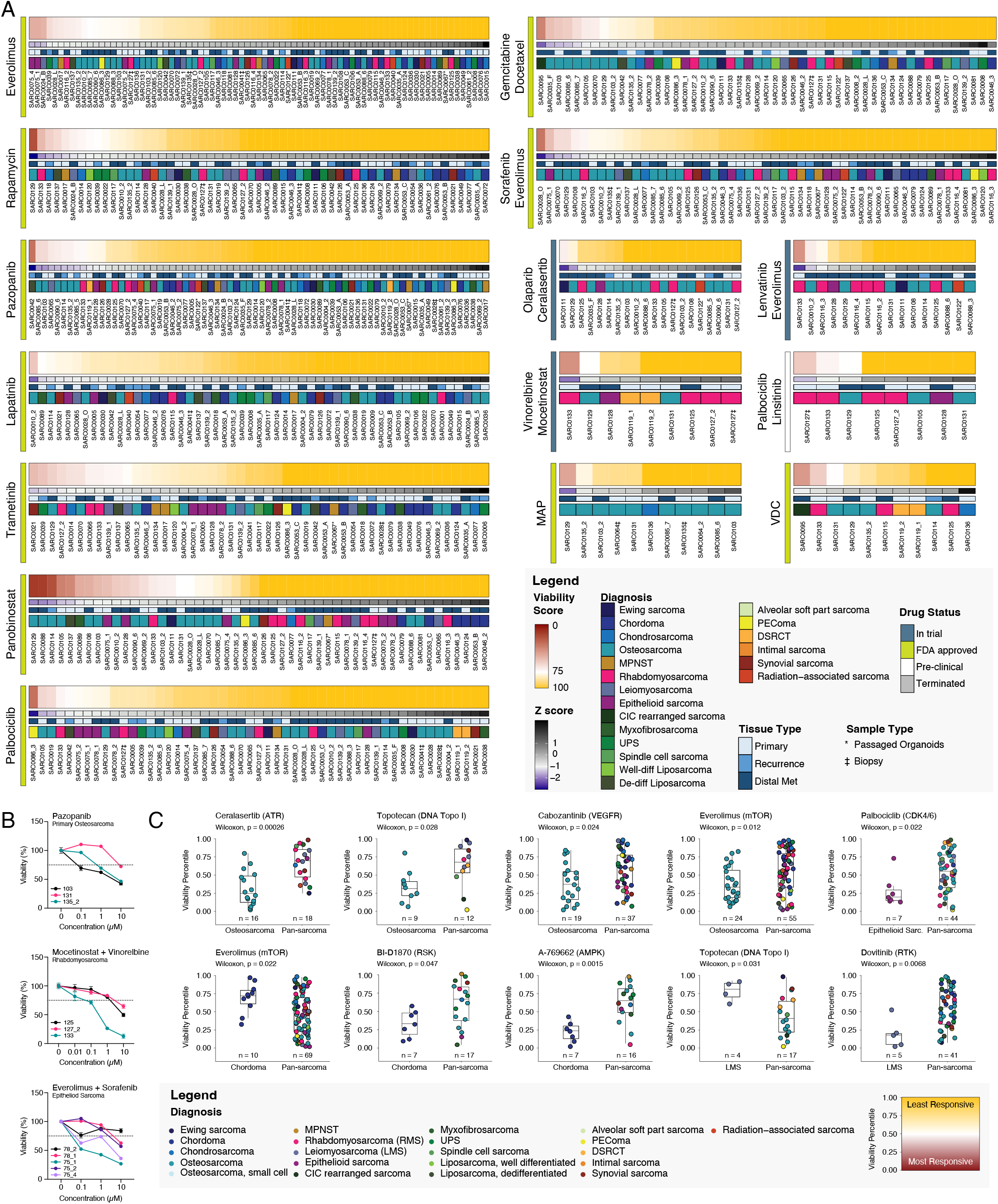
Sarcoma organoid sensitivity to treatment in high-throughput drug screening experiments shows a range of responses. (A) Heatmaps of organoid sensitivity to selected drugs of interest at 1 μM. The viability score represents each organoid model’s viability normalized to the mean response to treatment across all samples. Each column is a unique specimen, red indicates higher sensitivity to treatment than average. Colored bars underneath each heatmap represent the Z-score, lesion type, and diagnosis of each sample. *indicates passaged samples, denotes biopsies. (B) Dose-response curves of organoid viability when treated with selected therapeutic regimens. Percent viability is reported compared to vehicle-treated organoids for each individual sample. (C) Sensitivity rank plots comparing the response of organoids derived from the indicated diagnoses (left) against pan-sarcoma specimens (right). Samples are ranked from low residual viability percentile (most responsive samples) to highest residual viability percentile (least responsive samples). Primary drug targets are indicated next to each drug’s name. The color of each point represents the diagnosis of the individual samples.

### The landscape of drug sensitivity and resistance in sarcoma PDTOs

Sarcoma PDTOs were tested for sensitivity against chemotherapy or targeted drugs as single agents or in combination. We included therapies used for sarcoma in the clinic such as sorafenib and everolimus, gemcitabine and docetaxel, and methotrexate, doxorubicin, and cisplatin (MAP^4,5^, **Figure 3A and 3B**). To account for variations in drug efficacy, such as in case of molecules with known broad cytotoxic activities^37^, we calculated a viability score by normalizing the viability of each specimen to the average value observed across all samples treated with the same regimen (**Figure 3A**). By employing this approach, we identified individual samples that displayed sensitivities to specific therapies within our screened library. Sarcoma responses to a subset of single and combinatorial therapies is shown in **Figure 3**.

Among all the specimens screened against 10 or more compounds and included in the analysis (n=92), most samples had at least one significant hit (74/92, 80.4%), defined as ranking in the top 5% of viability scores for one or more of the regimens tested. Only 3 samples ranked in this category for a quarter or more of the compounds tested (3.3%), indicating broad and possibly nonspecific chemosensitivity. Conversely, 18 sarcomas did not rank in the top 5% for any compound (19.6%). This group of samples were generally screened against smaller drug libraries with an average of 54 regimens tested per sample vs 116 for samples with at least one hit (median: 25 vs 58 drugs or combinations), which indicates how smaller screens may be insufficient to identify effective regimens. Largely, samples showed responses to specific compounds. For instance, SARC0021, a synovial sarcoma, exhibited strong sensitivity to the MEK inhibitor trametinib^38^ (viability score: 54), yet ranked among the most resistant samples to pazopanib, palbociclib, rapamycin, and everolimus (**Figure 3A**). PDTOs derived from the SARC0086_3 PEComa had the highest sensitivity to palbociclib and showed strong resistance to pazopanib and to the combination therapy of sorafenib and everolimus (**Figure 3A**).

We also observed sensitivities to classes of similar compounds. Everolimus, a rapamycin analog and mTOR inhibitor, has demonstrated modest clinical efficacy in the treatment of various bone and soft tissue sarcomas^39^. To assess the sensitivity to everolimus within our sarcoma PDTO cohort, we tested a group of n=79 samples, yielding a range of responses (**Figure 3A**). Notably, among the 56 samples screened against both everolimus and rapamycin, 8 (14%) displayed positive responses to both drugs, ranking in the top quartile of responders. These included 3 osteosarcomas (SARC0129, SARC0010_2 and SARC0135_2), a de-differentiated liposarcoma (SARC0137) and a well-differentiated liposarcoma (SARC0120) as well as one MPNST (SARC0017) and an undifferentiated pleomorphic sarcoma (SARC0039). Conversely, organoids derived from the de-differentiated chondrosarcoma SARC0014 are the only example of remarkable sensitivity to rapamycin (top quartile of responders), yet resistance to everolimus (bottom quartile, **Figure 3A**). Some sarcoma subtypes, particularly bone tumors such as chondrosarcoma or chordoma, are considered broadly chemoresistant from a clinical standpoint^26,40,41^. This is consistent with our data, where these tumor types are often resistant to therapy (**Figure 3A**). As an example, we observed low response rates to everolimus for chordoma and chondrosarcoma PDTOs, accounting for 4/5 most resistant samples to treatment (**Figure 3A**). We observed similar patterns of resistance of chondrosarcoma and chordoma PDTOs to rapamycin and lapatinib (**Figure 3A**).

We also explored the effectiveness of combination therapies including regimens that are already in clinical use or in trials (**Figure 3A and 3B**). SARC0095, a CIC-rearranged Ewing-like sarcoma, exhibited significant sensitivity to gemcitabine and docetaxel (viability score: 36). Gemcitabine-based regimens have been reported as inducing two partial responses in a recent study investigating clinical management and outcomes for CIC-rearranged sarcomas^42^. The combination of vinorelbine and mocetinostat is currently in trials for rhabdomyosarcoma^43^. We observed an exceptional response for the RMS SARC0133 (**Figure 3A**), with heightened sensitivity across all tested concentrations (**Figure 3B**) when compared to the other three RMS samples that were evaluated for sensitivity to vinorelbine and mocetinostat. In summary, evaluation of the drug sensitivity and resistance profiles across a broad spectrum of sarcomas subtypes revealed largely specific and distinct patterns of drug response, underscoring the utility of the approach to pinpoint individual responders for more personalized and targeted therapeutic strategies.

### Intra-patient tumor heterogeneity in drug sensitivity

In the proportion of patients (21%, see above) for which multiple lesions were obtained, we have observed cases of concordant and discordant responses, even for metastases procured within the same surgery. Amongst the most sensitive organoids within our cohort were those derived from two different metastatic lung lesions of an epithelioid sarcoma patient with undifferentiated pleomorphic features, SARC0075_4 and SARC0075_1. These showed broad sensitivity profiles, both responding to 23% of the panel of drugs tested. For instance, their viability scores were the highest of the n=79 total samples screened against everolimus, indicating the highest sensitivity of all sarcomas tested. A third metastasis obtained during the same surgical procedure, SARC0075_2 had a different overall profile, responding to only 7.7% of all regimens and exhibiting marked resistance to everolimus across all doses tested (SARC0075_3, **Figure 3A**). We observe a similar behavior with sorafenib and everolimus administered in combination, with EC_50_s of 12.9 µM, 4.4 µM and 0.2 µM for samples 2, 4 and 1 respectively (**Figure 3B**). Considering all tested drugs, samples 1 and 4 were closely correlated, with a Pearson coefficient of 0.88. Sample 2 was the least similar to the other metastases, with Pearson coefficients of 0.70 and 0.72 for samples 1 and 4 respectively. Importantly, our data is consistent with clinical observations, as this patient exhibited notable clinical heterogeneity and a significant degree of diversity in tumor-to-tumor responses to various agents, including temozolomide. This highlights how the behavior measured ex vivo captures biological features of clinical relevance.

Repeat PDTOs established from a biopsy (SARC0127‡) and subsequent therapy-naïve tumor resection (SARC0127_2) for a case of rhabdomyosarcoma exhibited a shared lack of response to a number of regimens, including gemcitabine/docetaxel, olaparib/celarasertib and vinorelbine/mocetinostat combinations (**Figure 3A**). Despite a brief interval between biopsy and resection (18 days), we detected some distinct differences in sensitivities, including to the palbociclib/ linsitinib combination, yielding an overall Pearson coefficient of correlation of 0.79. Given the short interval between procedures and lack of neo-adjuvant treatment, the heterogeneity could be related to intra-tumoral variability and sampling bias^25^. In summary, our study pipeline can quantify differences in drug response within individual patients, emphasizing the importance of considering tumor-to-tumor and intra-tumoral heterogeneity when using any approach based on limited tissue sampling for selecting therapeutic strategies.

### Sarcoma PDTOs demonstrate subtype-specific responses to treatment

Though we detect a degree of heterogeneity in responses from patients sharing the same sarcoma diagnosis (**Figure 3A and 3B**), we have also been able to identify disease-specific trends. We compared the viability scores for each diagnosis against all others sarcomas using the Wilcoxon Rank Sum Test with Bonferroni adjustment for multiple comparisons correction (**Figure 3C and Figure S3**). We found osteosarcoma organoids to be significantly more sensitive than the pan-sarcoma average for the drugs ceralasertib, topotecan, cabozantanib, and everolimus, with p values of 0.00026, 0.028, 0.024, and 0.012 respectively (**Figure 3C**). Celarasertib is an ATR inhibitor currently in trial in combination with olaparib for osteosarcoma^44^.

In accordance with their well-documented chemoresistance to treatment^45,46^, chordomas were significantly less sensitive than the pan-sarcoma average to everolimus (p = 0.022, **Figure 3C**), alvocidib (p = 0.00046), apitolisib (p = 0.0071), and bortezomib (p = 0.000018). Yet, we identified 3/256 drugs for which chordoma PD-TOs established from more than two patients showed greater sensitivity than the broader population: a ribosomal S6 kinase (RSK) inhibitor, BI-D1870 (p = 0.047), an AMP-activated protein kinase (AMPK) activator, A-769662 (p = 0.0015, **Figure 3C**) and a EGFR/HER2 kinase inhibitor TAK-285 (p = 0.034). We observed a number of other trends in cohorts with smaller sample sizes such as chondrosarcoma PDTOs sensitivity to the FAK/Pyk2 inhibitor, TAE226 (p = 0.04), and to the PDK1 inhibitor, BX-912 (p = 0.038), RMS organoids to pazopanib (p = 0.049) and trametinib (p = 0.013), and UPS organoids to the IGF-1R inhibitor, BMS-754807 (p = 0.039, **Figure S3**). Lastly, leiomyosarcoma organoids were on average more sensitive than other sarcomas to the FGFR inhibitor dovitinib (p = 0.0068). However, they exhibited lower responses to topotecan (p = 0.031, **Figure 3C**), in line with previous clinical trial results showing that topotecan was ineffective in leiomyosarcomas^47^. These trends may aid in the identification of possible indications for therapeutic regimens.

### Clinical features associated with drug responses

We performed the same analysis described above to identify other correlates of response, including clinical features such as patient age at diagnosis, lesion type (primary, metastatic or recurrent disease), treatment history, and disease progression (**Figure 4**). We identified trends on small numbers of compounds yielding differences in viability scores for tumors exhibiting certain characteristics. For instance, tumors exposed to prior systemic therapies of any kind up to three months prior to tissue procurement appeared more sensitive than treatment-naïve sarcomas to molecules such as the mTOR inhibitor everolimus or the multi-TKI inhibitors cabozantinib, lenvatinib and cediranib (p=0.018, p=0.036, p=0.011, and p=0.017, respectively; **Figure 4 and Figure S4**) yet more sensitive to golvatinib (p = 0.036, **Figure S4**). Stratification by number of previous therapies (0 vs 1-2 vs 3+) highlighted an interesting trend: tumors exposed to more drugs tend to be more sensitive than naïve ones, such as in the case of pazopanib and danusertib on PDTOs established from patients exposed to three or more lines of systemic therapy (p=0.0062, and 0.011, respectively), or PHA−767491 and lenvatinib for sarcomas treated with one or two lines of systemic therapy (p=0.029, 0.035, respectively, **Figure 4 and Figure S4**).

**Figure 4.**
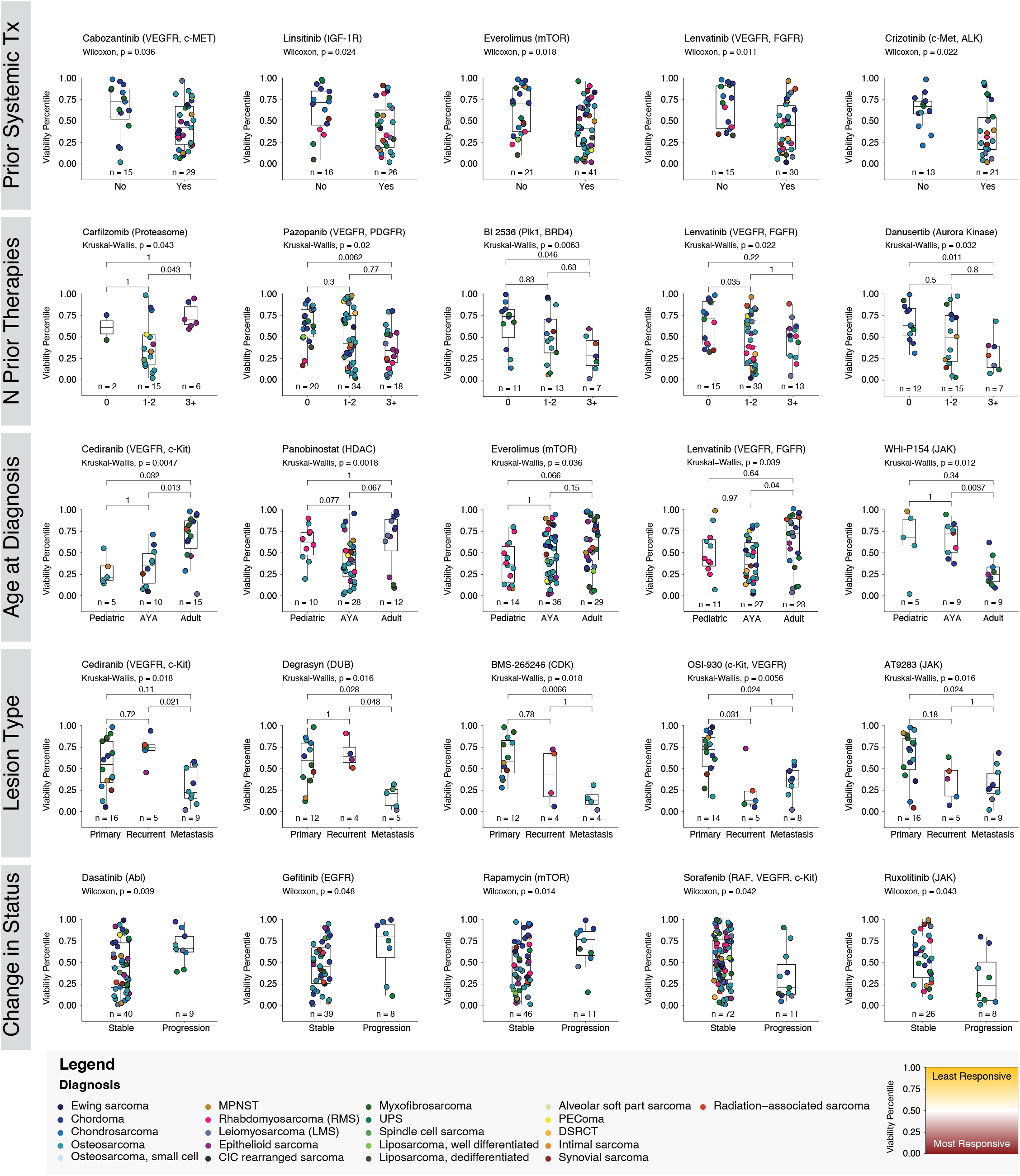
Sarcoma organoid sensitivity correlates with clinical attributes. Sarcoma samples are grouped by clinical features including patient age at diagnosis, lesion type, number of prior systemic therapies, prior systemic therapy within 3 months of sample procurement and change in disease status. All samples screened with the indicated drug are ranked from lowest viability percentile (reduced PDO viability, higher response to drugs) to highest viability percentile (increased PDO viability, lower response to drug) and plotted according to the rank. Primary drug targets are listed. The color of each point represents the specific diagnosis of the individual samples. Statistical significance is tested by performing a Kruskal-Wallis test with post-hoc Wilcoxon Rank Sum Test for pairwise comparisons with Bonferroni correction for comparisons across three classifications. For comparisons across two categories, a Wilcoxon Rank Sum Test was performed.

Patient age correlated with patterns of response; for instance, sarcoma PDTOs derived from adults showed greater resistance to cediranib compared to both pediatric (p=0.032) and AYA (p=0.013) sarcomas. PDTOs from adults were more sensitive than AYA to the JAK inhibitor WHI-P154 (p=0.0037, **Figure 4**). Taking into account the stage of disease, metastatic sarcoma PDTOs exhibited a tendency toward enhanced responses to degrasyn when compared to primary or recurrent tumors (p=0.028 and p=0.048) and to OSI-930 when compared to primary tumors (p=0.024, **Figure 4**).

Lastly, we investigated organoid response patterns with respect to changes in disease trajectory at time of follow-up. A change in status was defined as either a tumor recurrence following resection or the identification of metastatic disease from previously primary or localized recurrent tumors. PDTOs derived from patients who developed progressive disease were significantly more resistant to treatment with dasatinib (p=0.039), rapamycin (p=0.014), and gefinitib (p=0.048, Figure 4 **and Figure S4**). Tumors that rapidly progressed had increased sensitivity to thiazovivin, a ROCK inhibitor, as well as sorafenib and ruxolitinib (p=0.041, p=0.042, and p=0.043, respectively, Figure 4 **and Figure S4**). The observed differences in treatment response among various sarcomas were influenced by the age-specific prevalence of these tumors, as well as variations in disease management and treatment protocols^48–50^. Notably, cediranib is overall more effective on osteosarcomas than other histologies (**Figure S3**), thus it is not surprising that pediatric and AYA sarcoma PDTOs were more sensitive than adults, given the prevalence of osteosarcoma in our pediatric and AYA cohorts (Figure 4). Similarly, a large portion of metastatic samples in our dataset are osteosarcomas (17/40, 42%), while the chemoresistant chordomas are overrepresented among primary tumors (4/42, 10%). Therefore, while the trends reported are significant, they are strongly associated to specific sarcoma subtype characteristics.

### Target analysis of drug responses highlights vulnerable biological pathways in sarcoma

We conducted a comprehensive analysis of drug sensitivity profiles in sarcoma organoids to identify the molecular pathways contributing to drug responses (Figure 5 **and Figure S5**). We mapped each drug to their gene targets and pathways using the PubChem and WikiPathways^51^ (Figure 5) or KEGG pathways^52^ (**Figure S5**) databases. We then ranked the pathways from most impactful (Figure 5, red) to least impactful (Figure 5, blue) on the basis of their weighted components’ contribution to reduction in organoid viability scores (Figure 5). Interestingly, despite variations observed during the analysis of individual drugs (Figure 3), we consistently observed a tendency for samples derived from the same patient to cluster together. This pattern held true for SARC0103, SARC0078, SARC0075, SARC0139, and SARC0053, suggesting that sarcomas originating from the same patient share overarching vulnerabilities in molecular pathways, regardless of differences in tumor type, tumor location, timing of surgery, or response to specific drugs.

**Figure 5.**
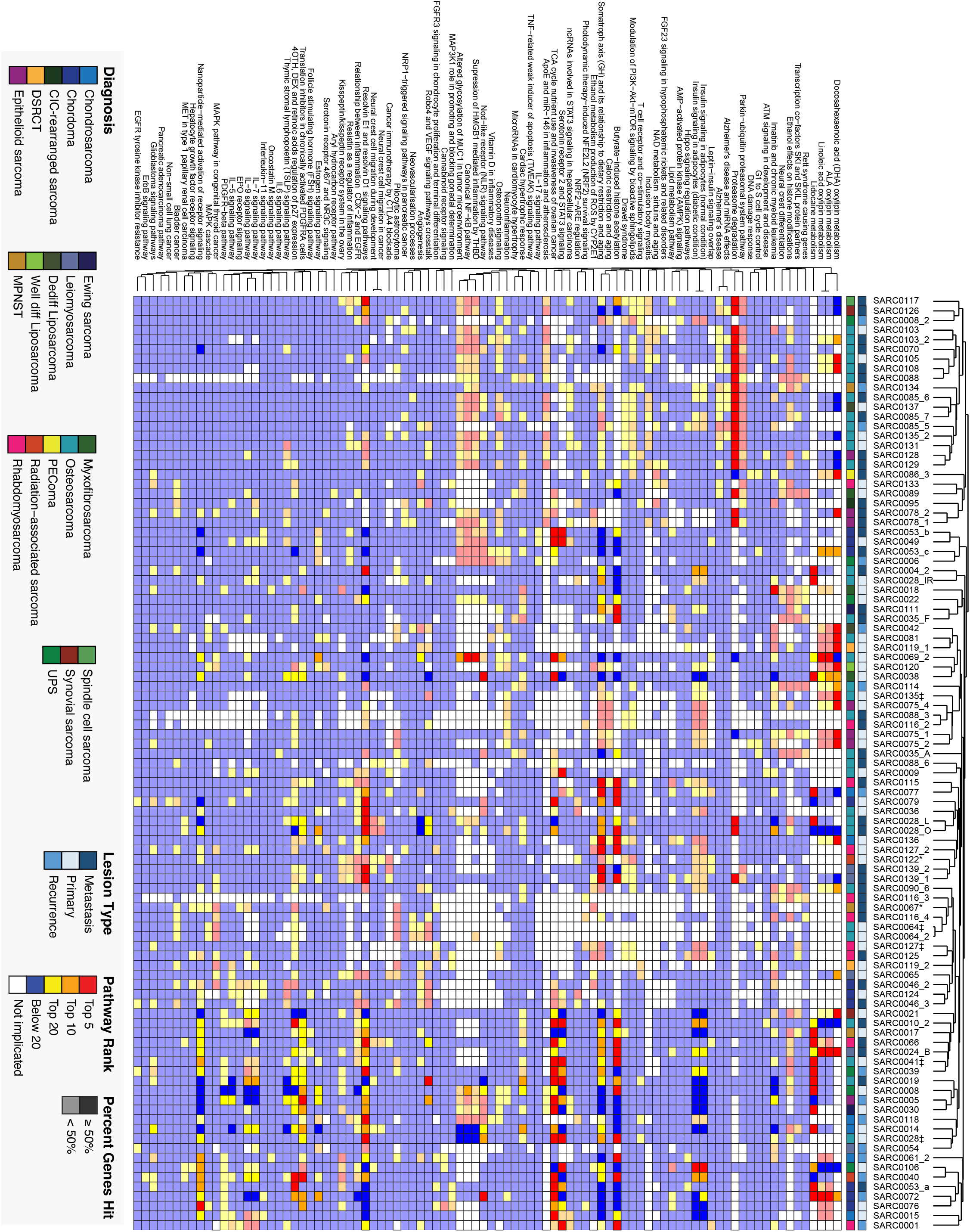
Landscape drug sensitivity patterns reveal vulnerable biological pathways. Heatmap showing the molecular pathways most sensitive to drug targeting for each screened sample from Wiki Pathways. Similar pathways are clustered together using the Jaccard distance and samples are clustered together by their Euclidian distance. Pathways are ranked independently for each sample based upon the results of drug screening experiments. Pathways targeted by the most effective drugs are ranked highest (red). Opaque squares indicate pathways in which more than 50% of the constituent genes were targeted in the drug panel. Pathways ranked in the top 50 for 20 or more samples are plotted. White squares indicate that the pathway was not targeted by any drugs in the screening experiments.

Sarcoma samples by and large did not cluster by histology, with some exceptions, highlighting the individual heterogeneity within sarcoma subtypes (Figure 5).

For instance, organoids derived from chordoma patients SARC0053 and SARC0049 clustered together due to strong sensitivity to treatments targeting genes in the serotonin receptor 2 and STAT3 signaling pathways as well as genes associated with TCA cycle (Figure 5), confirming our previous analysis^26^. These pathways contributed to the observed responses for a second group of samples encompassing both bone and soft tissue sarcomas (Figure 5). Perturbations in blood vessel-related pathways such as angiogenesis, Robo4 and VEGF signaling were linked to responses on a group of samples which included chordoma, DS-RCT, leiomyosarcoma cases (Figure 5). Additionally, we found a cluster of nine samples with vulnerabilities in the oxylipin metabolism pathway and, partially, in the insulin signaling pathway (Figure 5). Oxylipin derivatives have been linked to cancer cell proliferation^53^. Inflammation-related pathways were ranked as highly impactful across a broader set of 17 samples originated from a variety of histological subtypes (Figure 5). The same analysis performed using the KEGG pathways database allowed us to identify additional groups of pathways. For instance, EGFR and ErbB signaling pathways were significantly implicated in sensitivities observed in a group of samples which included chordomas (SARC0046_2 and 3, SARC0053_a, SARC0049, **Figure S5**), a subgroup of osteosarcoma samples (SARC0019, SARC0036, SARC0010_2, and SARC0041, **Figure S5**) and soft tissue sarcomas such as leiomyosarcoma and rhabdomyosarcoma (SARC0065, and SARC0127_2, respectively, **Figure S5**). Overall, this pathway analysis shows how PDTOs capture patient-specific features and susceptibilities, yet often transcend the histological subtype that currently dictates their treatment.

### Clinical availability of PDTO-selected drugs: a roadmap to actionability

As there is increasing interest in leveraging PDTO for precision medicine, we set to determine the degree of actionability of the drug sensitivities identified with our approach^26,37,54^. We define a drug or combination as actionable if it could be prescribed on- or off-label or accessed in the context of a clinical trial. To quantify actionability within our dataset, we generated a ranked list of the top five most effective regimens for each sample using the viability scores of single drugs and combinatorial treatments. We further refined the list by only including regimens in downstream analysis if the patient ranked within the top 10% of responders. This allowed us to eliminate generally ineffective drugs for highly chemoresistant samples. We then annotated the identified drugs by FDA status and recommendation on the basis of NCCN^4,5^ guidelines, if any (Figure 6).

**Figure 6.**
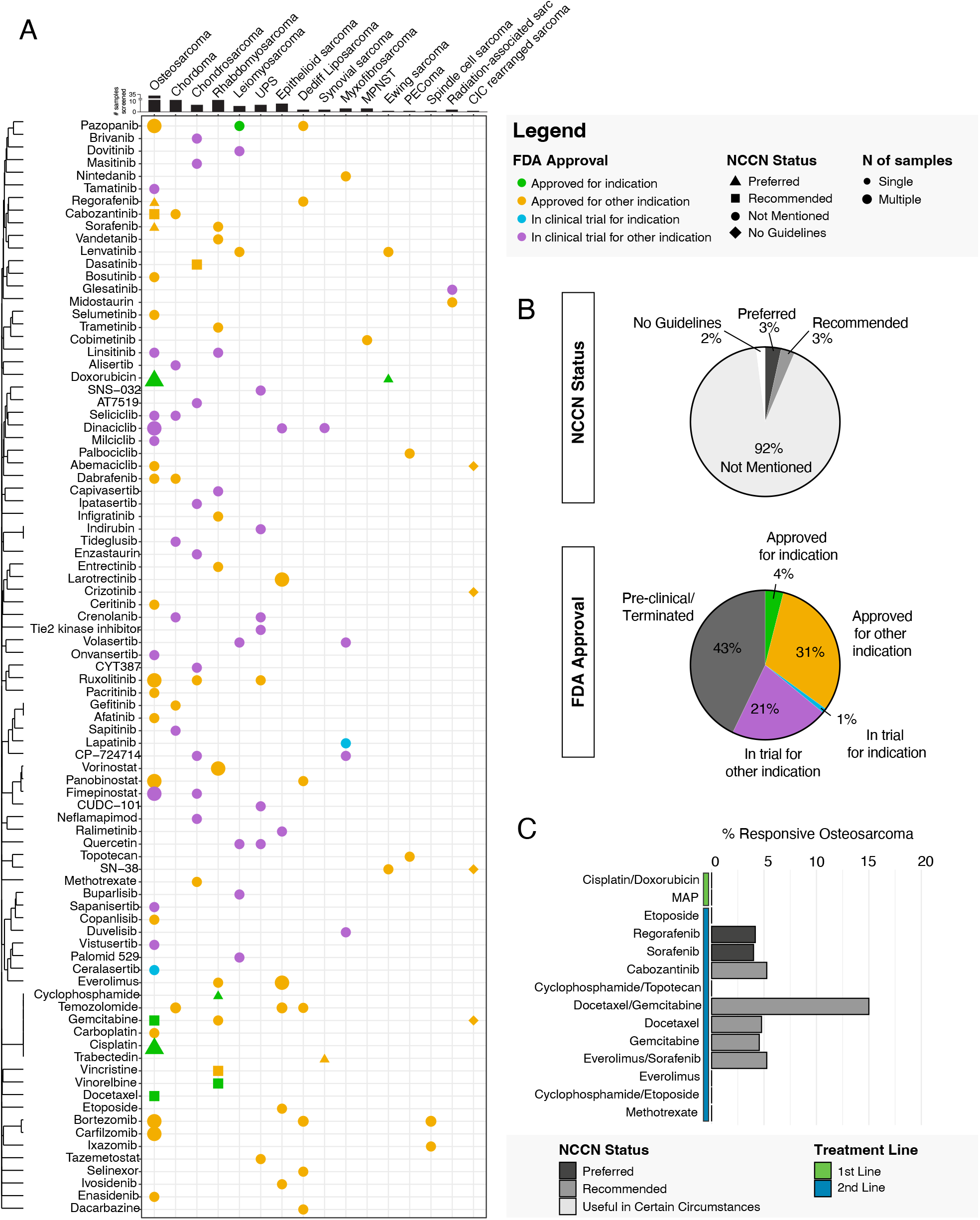
Actionability of PDO predictions determined by drug approval status. We selected drug-diagnosis pairs of interest by cross-referencing the five most effective drugs for each sample with the 10% most responsive samples for each drug. (A) Current FDA approval status and NCCN Guidelines recommendations are shown for each unique drug-diagnosis combination. Green and yellow colors indicate FDA approved drugs for the same cancer type or for other cancer indications respectively. Blue and purple represent drugs in trial for the same cancer or different cancers respectively. The shape of the symbols indicates NCCN Guideline status. Triangles shows drugs that are preferred, and squares recommended as per NCCN Guidelines. Circles are drugs not currently discussed in the guidelines4,5. Diamond shape signifies that the histologic subtype has no guidelines as of yet, such as in the case of DSRCT and CIC rearranged sarcoma. The size of the marker represents the number of samples for which each regimen is found amongst the most effective. Drugs are clustered by similarity in gene targets using Jaccard distance. The number of samples screened for each histologic subtype is shown in the bar charts above the graph. (B) Pie charts summarize the overall percentage of drugs that fall into each category for FDA approval and NCCN recommendations. (C) Percentage of responsive osteosarcoma PDOs to NCCN recommended treatment regimens.

After aggregating drugs by diagnosis, we identified n=203 drug-diagnosis pairs, that is, molecules found as effective in at least one patient with a certain subtype of sarcoma. This pairing allows us to verify approval status for specific indications (Figure 6A **and Figure S6**). Of these 203, 4% are FDA-approved for the same sarcoma type that was found as sensitive ex vivo, while 31% are approved for other cancers (Figure 6B). Given the rarity of sarcomas as a whole and limited number of existing trials, only 1% of the drugs identified are in clinical trials for the organoid-directed indication while 21% are in trial for other cancers (Figure 6B). Finally, 43% of the identified compounds are unavailable clinically either because they are in pre-clinical development or terminated (**Figure S6**), whether it be because of limited efficacy in an unstratified population, toxicities, or other factors^55^. Our analysis also identified n=24 combinatorial regimens fulfilling the criteria above (**Figure S6**). In aggregate, our approach identified at least one FDA approved or NCCN recommended regimen among the most effective for 59% (57/97) of sarcoma samples we screened. The majority of those samples had only FDA approved or NCCN recommended regimen (34/57), and only two samples had four effective drugs among those criteria (**Table S1C**). When looking at this data in the context of our sarcoma patients, we identified an effective approved or recommended therapy for 58% of patients screened (43/74), with 18 patients having only one approved therapy and only seven patients with four effective approved therapies (**Table S1A**). For patients from whom we collected multiple samples (12% of patients screened, 9/74), four patients had two approved drugs identified as effective, and five patients with four approved drugs (**Table S1B**).

We next took into account NCCN Guidelines for patients with bone^5^ and soft tissue^4^ sarcoma. The inclusion of a therapeutic regimen in these clinical guidelines requires substantial evidence of both safety and efficacy for each sarcoma subtype. Due to the small patient populations and highly heterogeneous disease, the vast majority (92%) of effective drug regimens we identified are not incorporated in the current guidelines. Only 5% of these are listed as preferred regimens by the NCCN for their corresponding histological subtype. These include etoposide, cisplatin, sorafenib, and regorafenib for osteosarcoma, cyclophosphamide for rhabdomyosarcoma, and doxorubicin for osteosarcoma and Ewing sarcoma (Figure 6A). An additional 3% of drugs are recommended for their respective diagnosis, including cabozantinib, gemcitabine, docetaxel, and everolimus for osteosarcoma, doxorubicin and vinorelbine for rhabdomyosarcoma (Figure 6A). The remaining 93% of drugs are beyond the recommendations of the NCCN Guidelines, either not mentioned (90%) or no guideline is available yet for the disease, as is the case for DSRCT and CIC-rearranged sarcomas. Our findings support the notion that functional precision medicine approaches may extend beyond the confines of current clinical guidelines to help match each patient to the optimal therapeutic regimens.

Osteosarcoma is a good test case for our approach; despite being a rare and heterogeneous tumor type, it has well established NCCN^5^ guidelines for both first and second line therapy (Figure 6C**, Figure S7, Table 1**). We categorized the drugs as listed in the guidelines as regimens used in first- or second-line therapy, as well as preferred, recommended, or useful in certain circumstances. Osteosarcoma is conventionally treated with a combination of methotrexate, doxorubicin, and cisplatin (MAP), in the first line setting^56^. While we observed sensitivities to MAP in osteosarcoma PDTOs, this was not among the top 5 most effective regimen in any of the samples tested (Figure 6C, **Table 1**). Of the second-line preferred regimens, the combination of gemcitabine and docetaxel produced effective responses in ∼15% (3/20) of the osteosarcoma PDTOs. Various second line treatments showed limited efficacy including regorafenib (1/24), cabozantinib (1/19), docetaxel (1/21), and gemcitabine (1/22, Figure 6C, **Table 1**). The broad heterogeneity in responses observed clinically is mirrored in our screening results. For instance, the combination of sorafenib and everolimus was effective in ∼5% (1/19), of screened osteosarcoma PDTOs including SARC0028_O (**Figure S7**). The limited responses are consistent with the clinical trial results in advanced and progressing osteosarcoma, with two partial responses (per RECIST criteria)^57^ and two minor responses (less than 30% and more than 10% reduction in lesion size per study criteria) recorded out of n=38 patients^58^.

**Table 1.** Response of osteosarcoma specimens to NCCN Guideline treatments. The responsive osteosarcoma samples meet the following criteria: the regimen was among the top 5 most effective regimens for a given osteosarcoma sample and the sample was among the 10% most responsive osteosarcomas to the listed regimen.

| Regimen | N responsive samples | N samples screened | Responsive Samples (%) | NCCN Status |
| --- | --- | --- | --- | --- |
| Cisplatin/Doxorubicin | 0 | 3 | 0 | Preferred |
| MAP | 0 | 10 | 0 | Preferred |
| Etoposide | 0 | 18 | 0 | Preferred |
| Regorafenib | 1 | 24 | 4 | Preferred |
| Sorafenib | 1 | 25 | 4 | Preferred |
| Cabozantinib | 1 | 19 | 5 | Recommended |
| Cyclophosphamide/Topotecan | 0 | 5 | 0 | Recommended |
| Docetaxel/Gemcitabine | 3 | 20 | 15 | Recommended |
| Docetaxel | 1 | 21 | 5 | Recommended |
| Gemcitabine | 1 | 22 | 4 | Recommended |
| Everolimus/Sorafenib | 1 | 19 | 5 | Recommended |
| Everolimus | 0 | 24 | 0 | Recommended |
| Cyclophosphamide/Etoposide | 0 | 1 | 0 | Useful in certain circumstances |
| Methotrexate | 0 | 15 | 0 | Useful in certain circumstances |

### Functional screenings provide orthogonal and complementary information to genomic sequencing

Both functional screenings and genomic sequencing can provide clues to tumor diagnosis and identify drivers of disease (Figure 7A and B). Mutations in the PIK3CA gene are linked to clinical responses to PI3K inhibitors such as alpelisib^59–61^. A tumor from a patient with metastatic undifferentiated spindle cell sarcoma (USS, SARC0117) was found to harbor the hotspot H1074L mutation in PIK3CA, associated with response to alpelisib^60,62,63^ on a clinical targeted sequencing panel. PDTOs from SARC0117 were among the strongest responders to alpelisib in our platform (Figure 7B). The metastasis we procured from SARC0117 had a concordant alteration in PIK3CA, as expected from the PDTO screening results (Figure 7A). Conversely, we obtained metastatic tissue from SARC0134, an MPNST that was reported to harbor the same PIK3CA hotspot mutation in the primary tumor. Nevertheless, SARC0134 PDTOs had no response to alpelisib (Figure 7B). Follow-up sequencing of the lesion demonstrated that no PIK3CA alteration was present in this metastasis (Figure 7A), confirming the screening results. Interestingly, the top responder to alpelisib, SARC0069_2, is a PDTO derived from a primary osteosarcoma that had no mutations in the PIK3CA gene found on sequencing (Figure 7A), which emphasizes how organoids may both validate sequencing results as well as identify biomarker-negative responders.

**Figure 7.**
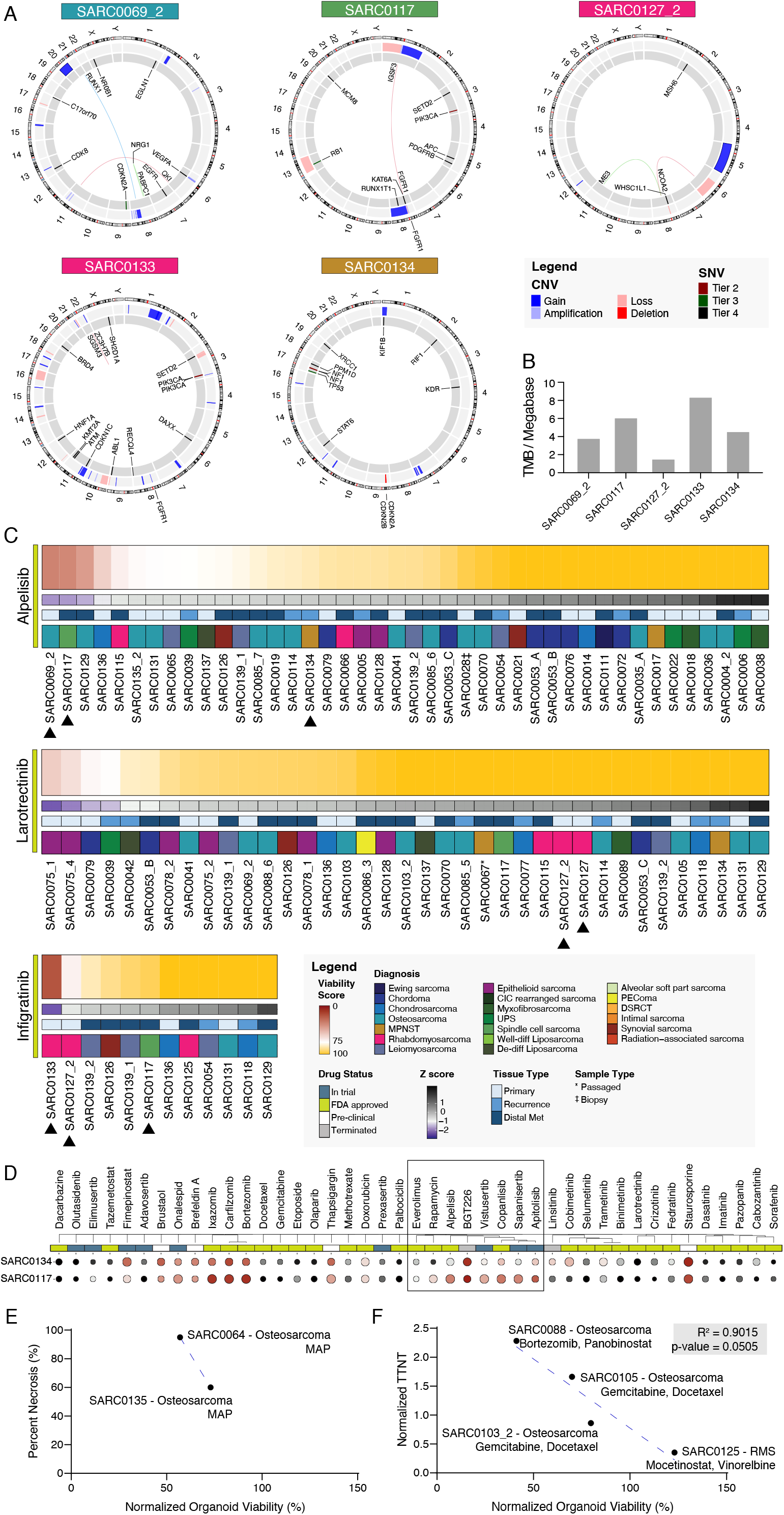
Organoids provide genomic and diagnostic information. (A) Summary of the genetic features of selected sarcomas. Sequencing was performed using OncoPanel. (B) Tumor mutational burden (TMB) for the same samples. (C) PDO viability scores for alpelisib, larotrectinib, and infigratinib. Black arrows indicate samples of interest. (D) Comparison of organoid sensitivity profiles for SARC0117 and SARC0134. The black box outlines PI3K targeting drugs with variable efficacy for samples SARC0117 and SARC0134. (C) Correlation between osteosarcoma biopsy response to MAP and percent necrosis clinically determined at time of tumor resection post neoadjuvant MAP treatment for the same patients. (F) Normalized time to next systemic therapy (TTNT) compared to PDO viability for therapeutic regimens screened on organoids and administered to the patient immediately following sample procurement using simple linear regression. TTNT of the matching therapeutic regimen is normalized to the TTNT of the regimen used immediately preceding specimen collection. TTNT greater than 1 indicates that the treatment of interest yielded longer TTNT compared to the previously administered treatment. Sample diagnosis and therapeutic regimen are annotated for each point.

The platform affords the opportunity to screen multiple drugs with shared targets to identify the most effective within a drug class. As an example, we tested eight drugs targeting the mTOR/AKT/PI3K pathway on SARC0117 (Figure 7). SARC0117 ranked among the top three most sensitive samples to apitolisib, alpelisib, copanlisib, BGT226, and vistusertib demonstrating its strong response across the entire class of mTOR/PI3K inhibitors (Figure 7C**, Figure S8**), in line with PIK3CA mutation driving the disease. However, in some cases we observe a distinct sensitivity to specific agents within a drug class. For instance, SARC0133, a rhabdomyosarcoma, showed the highest sensitivity to the FGFR targeting drug infigratinib relative to 11 other samples screened (Figure 7C). Sequencing revealed that SARC0133 harbors a FGFR1 gain on chromosome 8 contributing to its exceptional response to treatment (Figure 7A). Yet, there was minimal response to other FGFR-targeting molecules such as dovitinib (**Figure S8**), which exemplify the ability of functional screening to differentiate effective and ineffective drugs within the same class.

Due to the rapid results, our platform can be leveraged for diagnostic purposes for well-defined genomic alterations driving clear ex vivo and clinical responses to specific drugs. One such example is NTRK fusions driving responses to the FDA-approved NTRK inhibitor larotrectinib^64,65^. We received a portion of the diagnostic biopsy of a tumor, SARC0127‡, with clinical characteristics consistent with infantile fibrosarcoma. One of the hallmarks of infantile fibrosarcoma is the presence of NTRK fusions^64,65^. We established SARC0127‡ PDTOs and tested for sensitivity to larotrectinib. Results obtained less than a week from the diagnostic biopsy confirmed resistance to larotrectinib. Given the clear relationship between NTRK alterations and larotrectinib sensitivity both in vitro and in patients^66^, and overrepresentation of NTRK fusions in infantile fibrosarcoma, as 90% of infantile fibrosarcoma cases were found to harbor an ETV6-NTRK3 gene fusion^67,68^, we hypothesize that the tumor could be a different sarcoma subtype. Initial pathology review showed a high-grade sarcoma and FISH results that were available subsequent to our results showed the tumor was negative for ETV6 gene rearrangements, a classic finding in infantile fibrosarcoma. The tumor was sent for molecular testing and surgically resected, and 18 days after the original biopsy was diagnosed as a high-grade spindle cell/sclerosing rhabdomyosarcoma. PDTOs established from the resection (SARC0127_2) showed a similar pattern of drug sensitivity and resistance to the biopsy, including a comparable lack of response to larotrectinib (Figure 7). We validated the findings by performing FISH for NTRK 1, 2, and 3 fusions and confirmed that no abnormalities were present. Thus, our platform provided diagnostic clues within a week from the biopsy procedure, confirming the ability to rapidly yield information on the characteristics of each tumor.

### Evidence of correlation between PDTO and patient responses for matched treatment regimens

A major primary goal of developing personalized tumor organoid models is to leverage them to predict response to treatment. We have compared the PDTO screening results to clinical outcomes for two sets of samples. SARC0064‡ and SARC0135‡ are diagnostic biopsies from osteosarcoma treatment-naïve patients. Both patients received neo-adjuvant MAP between the time of biopsy and tumor resection. Upon resection, tumor tissue from SARC0064 was found to be 95% necrotic while SARC0135 had 60% necrosis. Tumor necrosis post neo-adjuvant therapy is correlated to osteosarcoma clinical outcomes, with necrosis >90% at resection indicative of long-term responses^69^. Ex vivo, SARC0064‡ PDTOs had a viability score of 57%, indicating a significant activity of MAP when compared to SARC0135‡ (viability score = 73, Figure 7E). Overall, the relative sensitivity of the organoids matched the clinical observations of necrosis percentage for these two patients (Figure 7E).

Furthermore, we identified a small cohort of patients who received a combinatorial treatment that was included in the library of compounds tested on the PDTOs. We included patients that received such regimen as next line after the sample procurement surgery. These include SARC0088, a patient with metastatic osteosarcoma treated with bortezomib and panobinostat, SARC0103_2 another metastatic osteosarcoma patient treated with gemcitabine and docetaxel, SARC0105 a primary osteosarcoma patient treated with gemcitabine and docetaxel, and SARC0125 a rhabdomyosarcoma patient treated with mocetinostat and vinorelbine. While the sample size is limited (n=4) we observed a correlation between normalized organoid viability and time to next treatment (TTNT) as a metric for real-world clinical response^70^. While further research with larger patient cohorts is warranted to validate these findings, the observed correlation, with a 95% confidence interval and an R^2^ value of 0.902 (p=0.05), supports the potential utility of PDTOs in predicting treatment response for sarcoma (Figure 7F).

## Discussion

Sarcomas are rare cancers, accounting for less than 1% of all diagnoses annually^3^, and encompass over a hundred subtypes. Some of these are indolent and hard to grow as cell lines or patient-derived xenografts^26^ while other subtypes are exceedingly rare and understudied^26,29^. Clinically relevant disease models that can be used to perform drug discovery studies are lacking for a vast majority of rare cancer subtypes^71^. We implemented a systematic approach to understand the landscape of drug sensitivity and resistance in sarcoma. Our study included patients at different stages of disease and with diverse treatment histories from 25 distinct histological subtypes of both bone and soft tissue sarcoma. The organoids accurately reflected the histological features of the original tumor. Sarcoma PDTOs have unique characteristics, with differences in shape, patterns of aggregation, and growth dynamics (Figure 2). Overall, this study demonstrates feasibility to rapidly generate personalized organoid models from diverse rare tumors.

Patient-derived organoids are of high interest for precision medicine due to their ability to recapitulate a patient’s response to therapy^23,72^. Feasibility of including organoid screening data in clinical decision-making workflows will depend on timing, reliability, and predictive power. We show here that it is possible to develop a translational pipeline for large scale functional precision medicine programs within a single institution. This entailed developing protocols for identifying suitable patients and retrieve clinical information, preserving tissue after surgery, and generating organoids for high throughput screening with results available in less than one week from procurement (Figure 1). By utilizing our automated pipeline, we generated and screened organoids from 124 samples including in total over 500 drugs and drug combinations. This allowed us to build a comprehensive database of drug sensitivity and resistance profiles^73^ (Figure 3).

The ability to screen a large number of drugs, particularly FDA-approved regimens or molecules currently in clinical trials, within the short timeline of our pipeline, could allow oncologists to obtain actionable information within a week of biopsy or surgery. Conventional approaches to precision medicine based on sequencing can pair a targetable alteration to an intervention for approximately 30% of patients^74^. We set out to quantify “actionability” of the sarcoma mini-ring organoid platform. We define actionable those cases for which one of the top five most effective drugs for which the patient is in the top 10% of responders is either FDA approved or in trials. Overall, our approach identified at least one drug that is FDA approved or in clinical trials for 59% of patients screened. This underscores the potential broad applicability of our functional precision medicine platform. Yet, while a large proportion of predictions includes an FDA approved molecule for an oncology indication, fewer than 5% were approved for same cancer the patient is diagnosed with and only 6% of drugs were listed in the NCCN Guidelines as preferred or recommended therapies (Figure 6B). As such, physician access to drugs will be a significant challenge in adopting functional precision medicine recommendations in the clinical setting.

Tumor organoids have been shown to be representative of clinical responses in a variety of carcinomas^29,72,75^. Ultimately, it is fundamental to determine the predictive power of sarcoma organoids. We observed concordance of organoid viability and clinical responses for two sets of samples. Firstly, organoid responses to MAP therapy for two treatment-naïve osteosarcoma biopsies (SARC0064 and SARC0135) correlated to necrosis assessed clinically post neo-adjuvant therapy (Figure 7E). Percent necrosis post neo-adjuvant therapy is an important prognostic factor in osteosarcoma, and correlates with long term outcomes^76^. We also attempted to determine possible correlations between organoid responses and time to next treatment (TTNT). In order to minimize confounding effects such as tumor evolution under treatment, we restricted our analysis to patients that went on to received one of the regimens included in our screening immediately after procurement surgery or biopsy. Despite the complexities of treatment and tumor heterogeneities, we could detect a trend with patients receiving a regimen that was effective ex vivo showing longer TTNT values than those whose organoids had measurable resistance (Figure 7F). This preliminary data is encouraging and justifies extending this analysis in the context of a clinical trial. The overall goal of any precision medicine approaches, including one that is organoid-based, is to provide valuable information to identify effective treatments, avoid ineffective therapies and ultimately improve overall outcomes. If coupled with genomic analysis, such assays can also help to identify the most effective molecule within a drug class (Figure 7 **and Figure S8**). Perhaps the biggest hurdle to precision medicine remains sampling bias. This is particularly relevant for highly heterogeneous sarcomas and advanced, disseminated disease. PDTOs captured both the inter-patient heterogeneity observed in clinical trials of sarcoma (Figure 3) as well as intra-patient heterogeneity. For instance, we observed differential sensitivity to the multi-receptor tyrosine kinase (RTK) inhibitor pazopanib in osteosarcoma organoids (Figure 3B). Clinically, pazopanib has shown partial response or stable disease in 13 (68%) out of 19 metastatic osteosarcoma patients, with no biomarker available to stratify sensitive and resistant cases^77,78^. We found marked intra-patient heterogeneity in several patients, including cases with synchronous or metachronous lung metastases. For instance, we observed marked differential sensitivities in synchronous lesions from an epithelioid sarcoma case (SARC0075, Figure 3B), as well as in two osteosarcoma metastases from patient SARC0028 (SARC0028_O and SARC0028_L, Figure 3).

The utility of landscape analysis of sarcoma drug response extends beyond precision medicine applications. Our data emphasized unexplored relationships between drug sensitivity and relevant clinical attributes such as histology, patient age, type of lesion, and treatment history (Figure 4). We found patterns of resistance to specific drugs such as the mTOR targeting molecule rapamycin (Figure 4) or sensitivities to sorafenib and ruxolitinib (Figure 4) that were shared between samples obtained from patients who had rapidly progressed over the course of this study. The identification resistance or sensitivity patterns associated with clinical attributes will help to delineate biomarkers for bone and soft tissue sarcomas.

## Methods

### Patient Sample Collection

The protocol for collecting and processing tumor tissue has been previously described^9,18^. In summary, fresh tumor specimens are obtained from consenting patients (UCLA IRB #10-001857, 11-003254, 19-002214). Solid tumors are minced and digested with collagenase IV (200 U/mL) to yield a suspension of single cells/small clusters. The cells are then transferred to a new tube, followed by red blood cell lysis with Ammonium Chloride Solution (Stem Cell Technology). Cells are then strained using a 100µm filter before counting and viability assessment using a Cellometer Auto 2000 (Nexcelom).

### Organoid Generation

Primary cells are resuspended in a 3:4 solution of Mammocult medium (Stem Cell Technology) and Matrigel (Corning)^17,26,29^. The mixture is kept on ice throughout the organoid seeding process to prevent premature crosslinking. We seed the organoids for drug screening by distributing 10µL of solution around the perimeter of the bottom of each well of a 96-well plate. We incubate the material for 30 minutes at 37°C to solidify the gel before adding 100µL of Mammocult medium to each well. Organoids cultured for histological and molecular analyses are seeded in 24-well plates. We seed 100,000 cells in 70µL of the Mammocult-Matrigel solution around the perimeter of each well of a 24-well plate. Each well plate is imaged using a high-content microscope (Celigo, Nexcelom) every 24 hours.

### Drug Screening

After allowing the organoids to grow and develop for 3 days, we perform drug treatments using a panel of targeted agents and chemotherapies to assess sensitivity following our published protocols^9,18^. First, we remove the medium from each well using an automated fluid handler (Microlab NIMBUS, Hamilton or epMotion 96, Eppendorf) and replace it with pre-warmed Mammocult containing the desired drug concentration and 1% DMSO. Each plate contains its own positive and negative controls for normalization. The positive control is 10 µM staurosporine and the negative control is 1% DMSO. Organoids are incubated at 37°C and 5% CO_2_ throughout drug treatment. After 24 hours, we exchange the medium with fresh, drug-loaded medium. After 2 days of treatment total, organoid viability is assessed with an ATP assay (CellTiter-Glo, Promega). The organoids are released from the matrix with dispase followed by addition of the CellTiter-Glo reagent. After 30 minutes, luminescence is measured using a SpectraMax iD3 plate reader (Molecular Devices).

### Database

We maintain a PostgreSQL relational database that stores coded non-identifiable patient and sample information of our biobank. We implemented several external databases to our database such as gene pathway data from WikiPathways^51^, and mechanistic targets of our drug library from PubChem^79^ and literature. After organoid plates undergo drug screening, we use a Python-based, custom XML parser to upload the luminescence data to the database. This data is then connected to a sample collection that contains additional information about the patient and sample procurement using Django, a Python-based web framework. The drug treatment used for each well is also manually uploaded to the database. All downstream analysis is performed using R (v4.2) and begins by querying the database.

### Drug Screening Analysis

We screened each drug with either n = 1 or 2 at a concentration of 1 μM, with the exceptions of platinum agents, cisplatin and carboplatin, which we screened at 25 and/or 50 μM. Plate-level statistics including the Z’-factor^80^, and robust Z’-factor^36^, are calculated for each plate of organoids and are used as inclusion metrics for subsequent analysis. The thresholds for inclusion in this study are the following: Z’-factor > 0.2 or robust Z’-factor > 0.2. These criteria were selected to exclude plates that have an insufficient statistical effect size which are prone to false positives and negatives in high-throughput assays. We also excluded plates of experiments from passages or thawed samples if they were performed for the purpose of re-screening except for one sample where we included both the fresh and thawed experiment plates due to low QC metrics from the fresh experiment. We also excluded one sample from the analysis due to it being a benign tumor. We used the luminescence values of staurosporine screened at 10 μM as a positive control and substituted with 1 μM when not available. For plates that were screened with only one well of staurosporine 1 μM, the values were pooled across the plates as a positive control for the experiment. For plates that were included in the analysis, the luminescence measurements from the ATP assay are normalized to the mean luminescence of the negative control (1% DMSO) wells to calculate percent viability. For each drug treatment, the viability of each sample is normalized to the mean response of all samples treated with the drug of interest to calculate the viability score. Response rank percentile is calculated by dividing the rank of each sample and dividing it by the total number of samples screened with each drug.

### Quantification of samples sensitivities

We quantified the total number of drugs screened on each sample that are included in the analysis and filtered to only include samples screened with 10 or more compounds. We then filtered the data for all the drugs we have screened on a minimum of 3 samples to include only samples that rank in the top 5% of viability scores for one or more of the regimens tested to count the number of sensitive drugs identified per sample among the 5% cutoff. We then normalized the number of sensitive drugs identified for each sample by dividing over the number of total drugs screened on the sample to account for differences in the size of the drug library screened on each sample. We then quantified the number of samples with broad sensitivity by having a ranking of over 25% sensitive drugs or drug combinations identified.

### Histopathology

Histopathology analysis is performed on the tissue of origin and the organoids derived from the collected specimens. Organoids are prepared for histopathology after 5 days of culture. Each well is washed with 1 mL of phosphate buffered saline (PBS) prior to fixation with 500 µL of 10% buffered formalin (VWR, 89379-094). After at least 24 hours of fixation, the organoids are removed from the 24-well plate and transferred to a conical tube. They are washed with PBS prior to the addition of 5 µL of Histogel (Thermo Fisher Scientific, HG-40000-012). The sample in Histogel is then transferred to a cassette and paraffin embedded. We section organoid blocks at 8 µm and mount them on Superfrost Plus Microscope Slides (12-550-15, Fisher Scientific). Hematoxylin and eosin staining was performed on the parent tumor and the resulting organoids derived from tissue collection according to standard protocols. All images were acquired using the Revolve Upright and Inverted Microscope System (Echo Laboratories).

### Growth Quantification with Image Analysis

We image the organoids daily using a high-content microscope that scans two focal planes per well. The resulting whole-well images are exported in TIF format at a resolution of 1 µm/pixel. We then implement our previously developed methodology to segment and quantify regions of the image containing organoids^26^. We use a convolutional neural network with a U-Net architecture^34^ to segment the regions of the images containing organoids. This model is based on a Res-Net-34 model^81^ trained on 223 manually-labelled images spanning an array of tumors of origin to capture the diverse organoid morphologies observed in this study. In the manually-labelled dataset, only in-focus organoids are marked for inclusion in the area calculation; this is done to minimize measuring the same organoid across both focal planes. The original weights were derived from a model pretrained on the ImageNet dataset^82^ and the final model was trained over 80 epochs using a cross-entropy loss function. The trained model was then used to segment each image by splitting the image into 512×512 pixel sections, applying the model to each section, and reassembling the sections to recreate the whole segmented image (16,896×16896 pixels). We then implement OpenCV to calculate the total area of the organoids in each focal plane. This area is then averaged across both focal planes and growth is measured by normalizing to the area covered by organoids on the first day of imaging of the same well. The resulting data was then plotted using GraphPad Prism as normalized area over time.

### Targeted Sequencing

Select samples are sent to the Center for Advanced Molecular Diagnostics (CAMD) at Brigham and Women’s Hospital for analysis using the OncoPanel v3^83^. Sequencing was performed on pre-sectioned 10 slides from non-decalcified FFPE tissue blocks. Slides are then shipped to CAMD for sequencing and analysis. We plotted the results in a circos plot using R.

### Fluorescence In Situ Hybridization (FISH)

Identification of NTRK 1, 2, or 3 fusions was performed by preforming Fluorescence In Situ Hybridization (FISH). Pre-sectioned slides from non-decalcified FFPE tissue blocks from select samples are sent to NeoGenomics for performing their NTRK 1, 2, 3 FISH Panel (88374×3).

### Pathway Analysis

Protein targets for each drug in our library were annotated and obtained from PubChem^79^ and literature. We selected only targets that are within 10-fold of the second-lowest reported value for Kd (dissociation constant) or IC50 (median inhibitory concentration) among the targets. To perform the pathway analysis, we mapped each drug and their protein target with values of 0 and 1, with 1 indicating that a drug targets a protein, and 0 indicating the lack of protein among the drug targets. We filled the values in a matrix composed of n_drugs_ × m_protein_. To adjust for the degree of impact of protein targets on the viability of samples, we multiplied each row in the matrix by a weight proportional to the mean viability of the organoids treated by each drug [1 − (mean viability/100)]. We then multiplied this matrix by a vector of 1s to obtain a row-wise summation of the protein target viability values. To adjust for protein targets that are targeted by multiple drugs in our library compared to targets associated with drugs that are less represented in our panels, we normalized the row vector by dividing each element by the sum of non-zero column entries in the n_drugs_ × m_protein_ matrix. We then mapped this list of proteins to the canonical pathways defined by the WikiPathways Database [version 20220710]^51^. We excluded pathways that are not biologically pertinent to cancer, such pathways related to microorganisms and pathogens, as well as the newly added coronavirus disease (COVID)–related pathways. We populated a new matrix comprised of n_pathway_ x m_protein_, with 1s indicating the presence of a protein in a particular pathway or 0s indicating its absence. We then normalized the rows in the pathway matrix to account for the differences in the number of proteins included in each pathway. To obtain the relative effect that targeting a specific pathway has on the viability of sarcoma organoids, we multiplied the n_pathway_ × m_protein_ mapping matrix by the normalized n_drugs_ × m_protein_ vector. The resulting vector represents the relative impact that targeting a given pathway has on the viability of the organoids. We then ranked the scored pathways for each sample to compare the impact of each pathway on a viability of a sample between organoids of different sarcoma subtypes.

### Assessment of Drug Availability

Based on our drug screening data, we created a list of the five most effective therapeutic agents for each sample screened. We then created a second list of the top 10% most responsive samples to each drug. We considered only sample-drug pairs that appeared in both lists for further analysis and mapped each sample to its diagnostic subtype. For each drug-diagnosis pair, we manually annotated the inclusion of each therapy in the NCCN Guidelines^4,5^ as well as the current FDA approval status for each drug with respect to the histological subtype.

## Acknowledgments

We would like to acknowledge the Translational Pathology Core Laboratory at UCLA for assistance performing immunohistochemistry studies and the CAMD core at Dana-Farber Cancer Institute for assistance in targeted sequencing of tissue. We would also like to thank R. Damoiseaux and the UCLA MSSR Core for providing part of the drug libraries included in this study. We would like to thank the UCLA JCCC BASE team for their consultation on our statistical analysis. We would like to acknowledge Dr. Brian Kadera and Dr. Brooke Crawford for their involvement in consenting patients and collecting tissue for analysis and Matthew Mapua for technical support in initial studies. We are grateful to all the patients and patient families that participated in this study. This work was supported by the NCI grants R01CA244729 and R01CA244729-03S1 (to AS and PCB), a Slifka Foundation Award (to AS and NF), a UCLA JCCC Program Leader Vision Award (to SN), and a UCLA DGSOM Seed Award (to AS, PCB, NF, AK and JY). Additional funding was provided by a JCCC Fellowship Award (to PT), and NIH U24CA248265 (to PCB).

## Supplementary Material

### Supplementary Tables

**Table S1.**
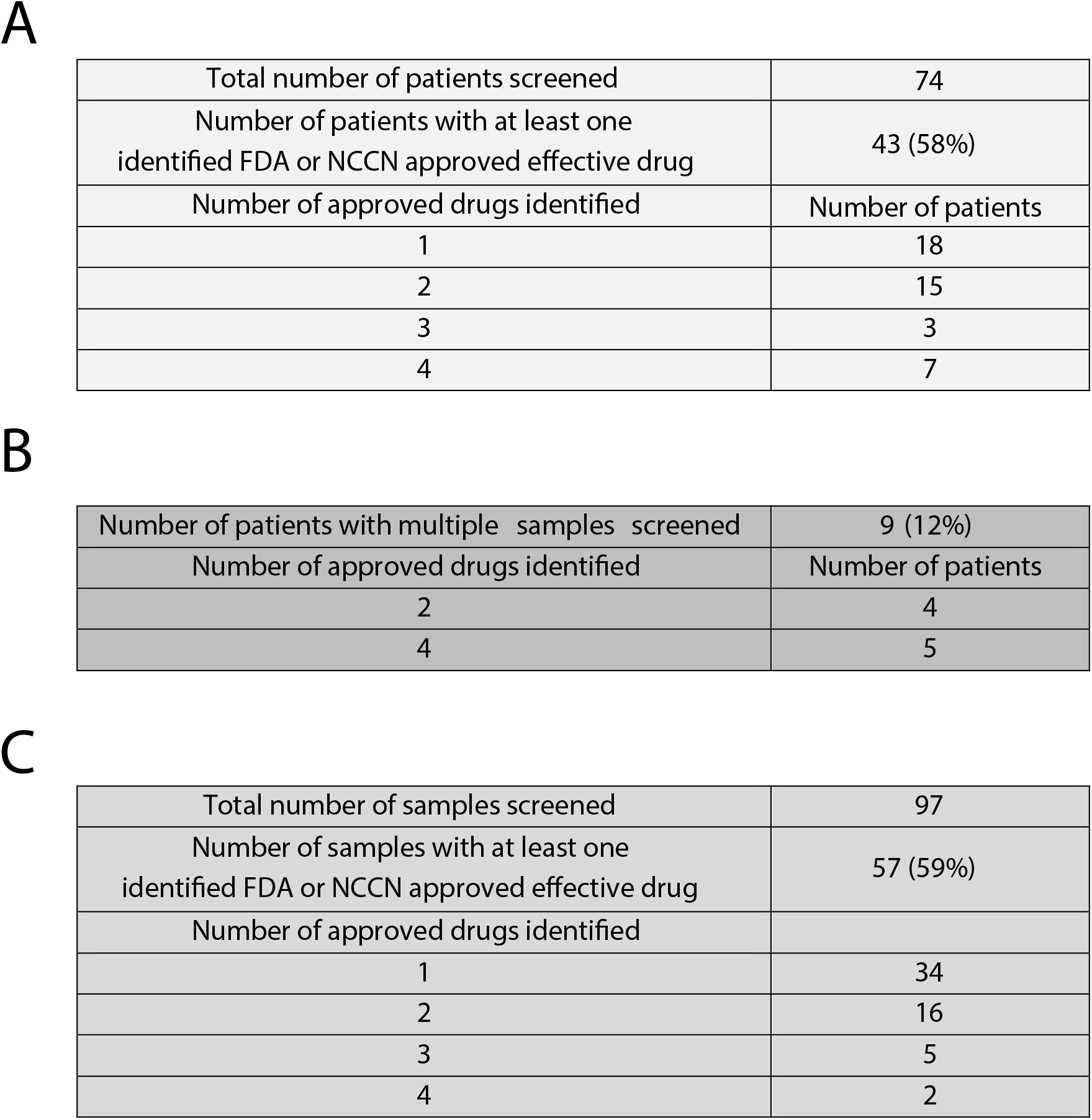
Breakdown of number of effective drugs identified that are FDA approved or NCCN recommended or preferred by patients (A), patients with multiple samples procured (B), and samples (C).

|  |  |
| --- | --- |
| Total number of patients screened | 74 |
| Number of patients with at least one identified FDA or NCCN approved effective drug | 43 (58%) |
| Number of approved drugs identified | Number of patients |
| 1 | 18 |
| 2 | 15 |
| 3 | 3 |
| 4 | 7 |
| B |  |
| Number of patients with multiple samples screened | 9 (12%) |
| Number of approved drugs identified | Number of patients |
| 2 | 4 |
| 4 | 5 |

|  |  |
| --- | --- |
| Total number of samples screened | 97 |
| Number of samples with at least one identified FDA or NCCN approved effective drug | 57 (59%) |
| Number of approved drugs identified |  |
| 1 | 34 |
| 2 | 16 |
| 3 | 5 |
| 4 | 2 |

### Supplementary Figures

**Figure S1:**
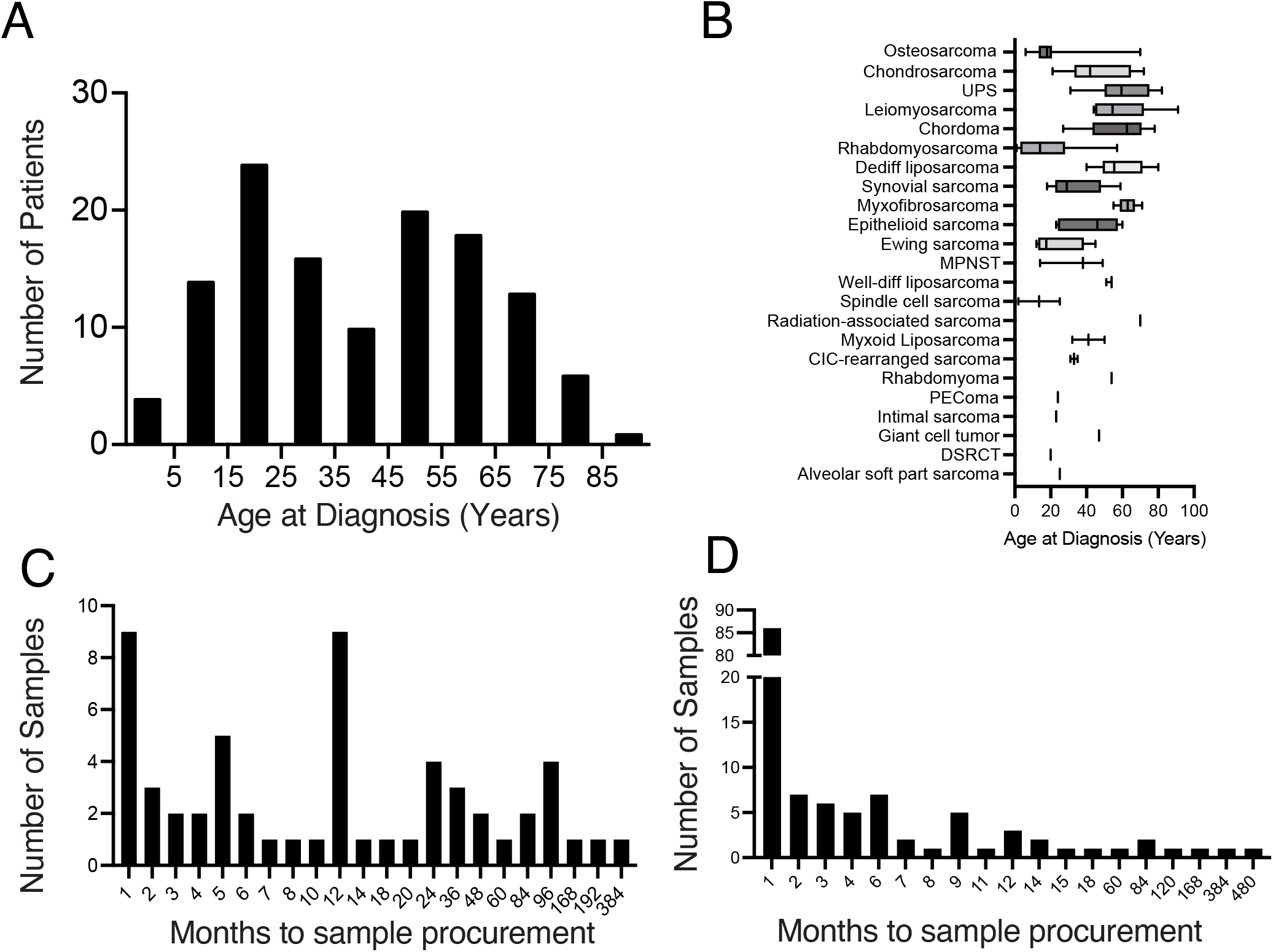
Patient and sample characteristics. (A) Age at diagnosis for the sarcoma patient cohort. (B) Age distribution by subtype. (C) Distribution of time to sample procurement in months from prior exposure to radiation therapy and (D) prior exposure to systemic therapy for our sarcoma tumor samples.

**Figure S2:**
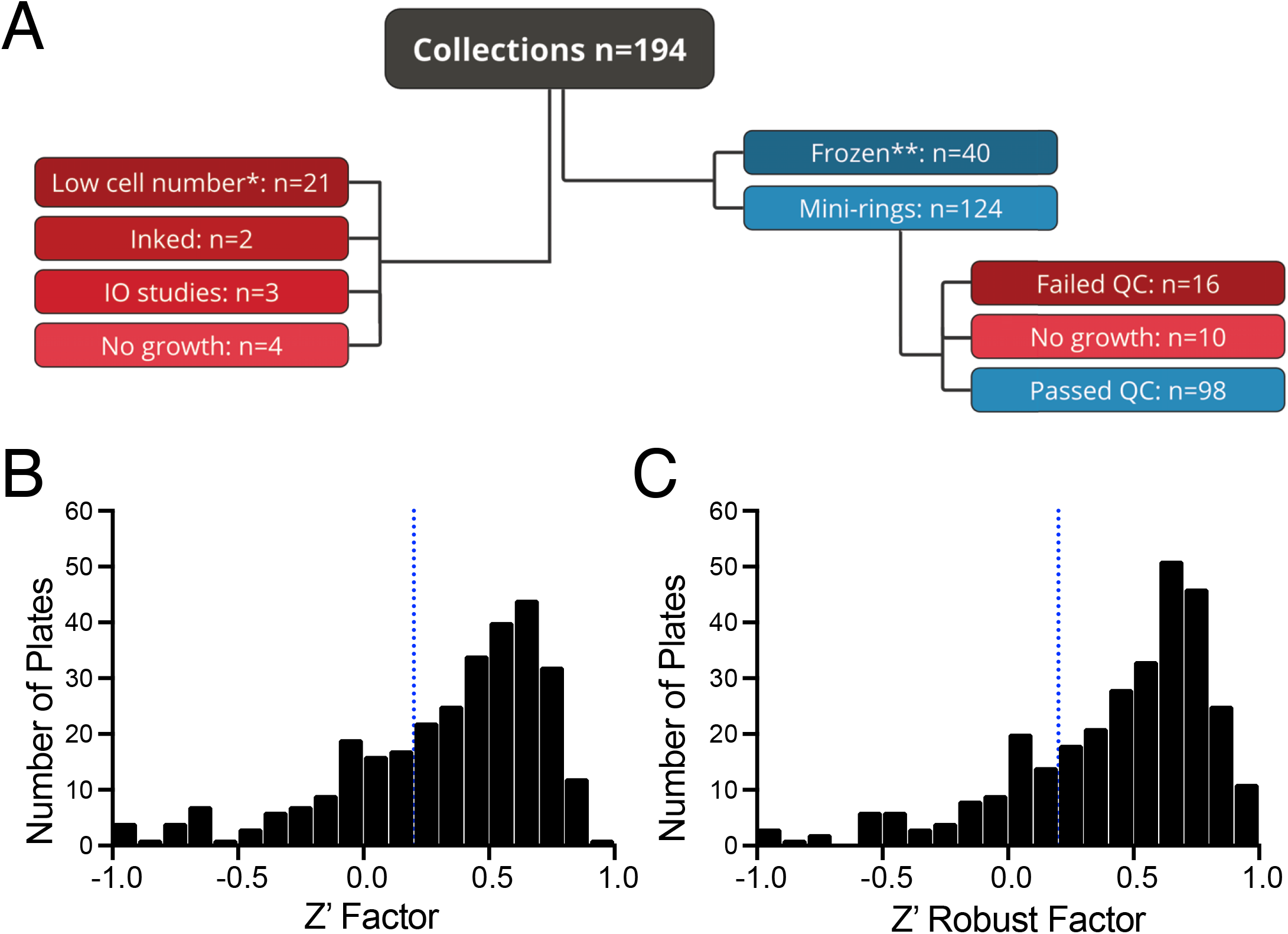
Sample pipeline and assay characteristics. (A) Breakdown of all sarcoma samples procured and yield of successful sarcoma organoid establishment, * = less than 250K viable cells harvested, ** = 48% due to covid restrictions in operations. (B + C) Distribution of Z’ factor and robust Z’ factor of the screened sarcoma plates. Dotted vertical line indicates cutoff of 0.2 for both Z’ factor and robust Z’ factor.

**Figure S3:**
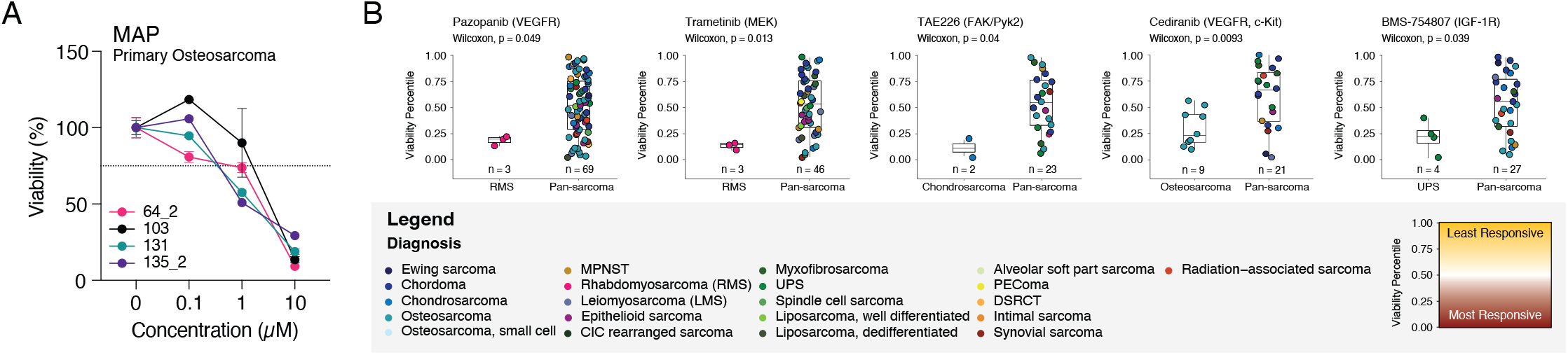
Dose response curves and diagnosis correlations. (A) Dose response curves of MAP screened on osteosarcoma PD-TOs. (B) Additional drugs with correlation between specific sarcoma diagnosis subtypes compared to pan-sarcoma samples. All samples screened with a drug are ranked from lowest viability (low viability percentile) to highest viability (high viability percentile) and plotted according to the rank. The color of each point represents the diagnosis of the individual samples screened with the drug of interest. Statistical significance is tested by performing a Wilcoxon Rank Sum Test with Bonferroni correction.

**Figure S4:**
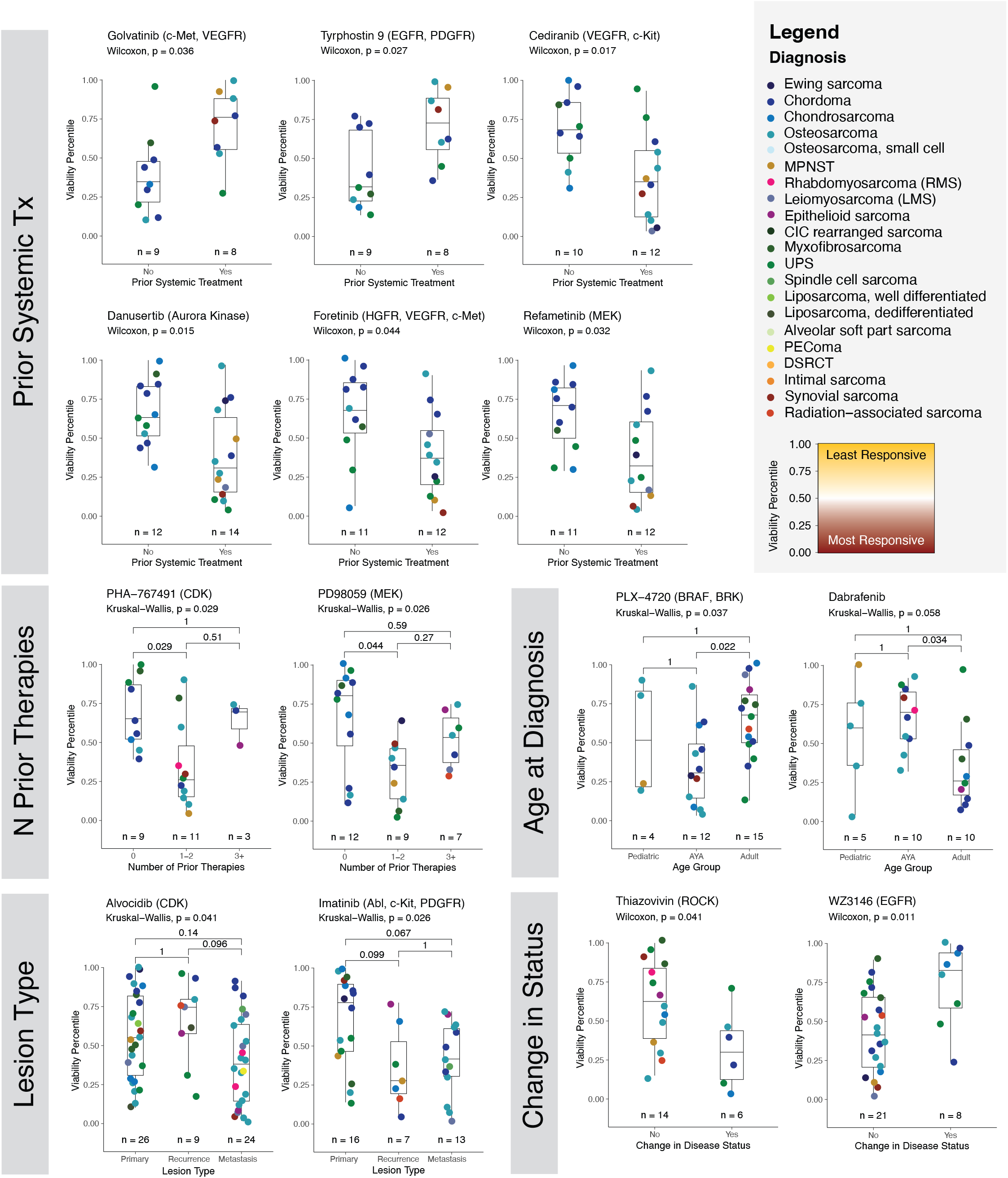
Additional drug with correlation between organoid sensitivity and prior systemic therapy within 3 months of tissue procurement, number of prior systemic therapies, lesion type, age at diagnosis, and progression of disease. All samples screened with a drug are ranked from lowest viability (low viability percentile) to highest viability (high viability percentile) and plotted according to the rank. The color of each point represents the diagnosis of the individual samples screened with the drug of interest. Statistical significance is tested by performing a Kruskal-Wallis test with post-hoc Wilcoxon Rank Sum Test for pairwise comparisons with Bonferroni correction.

**Figure S5:**
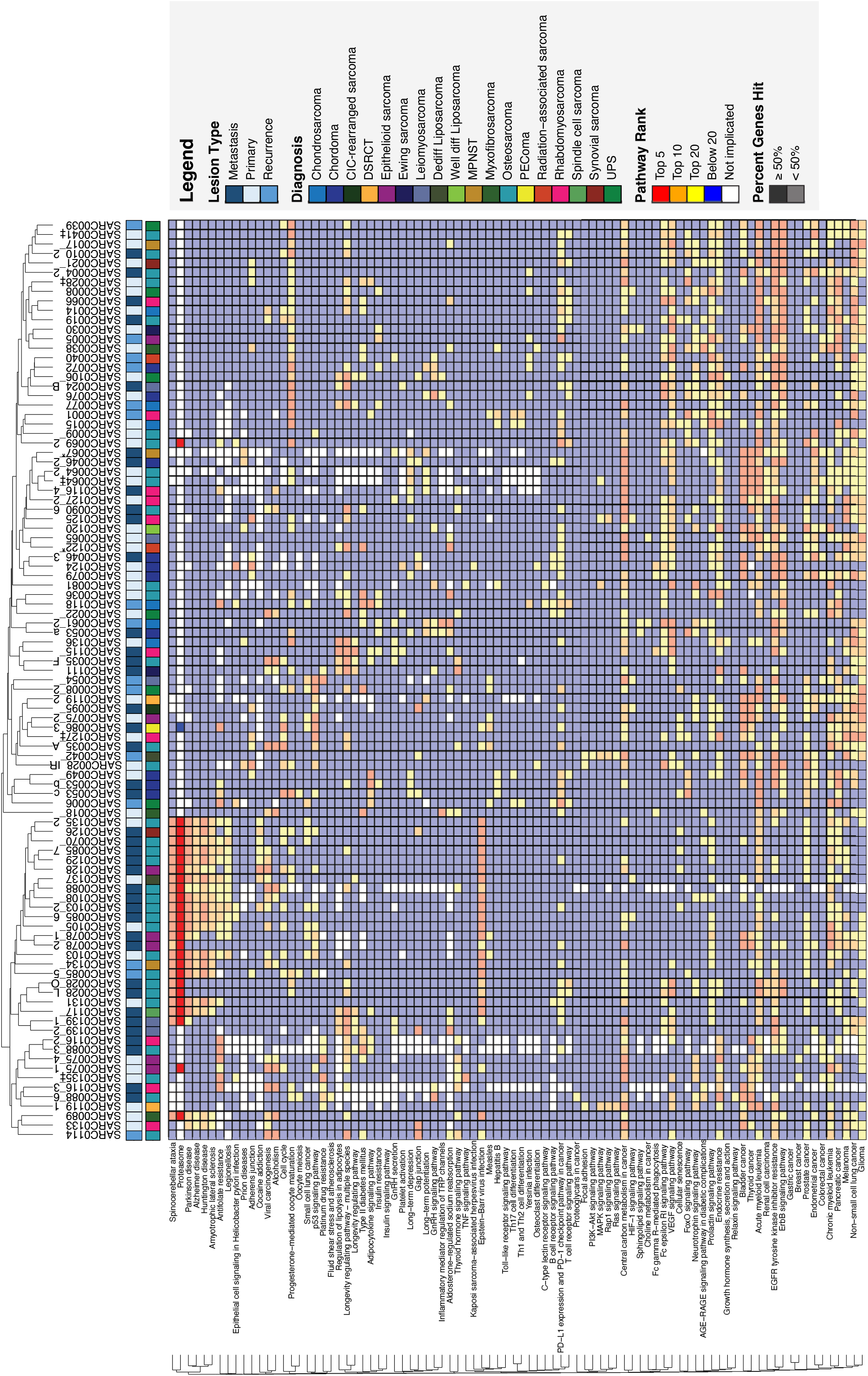
Additional landscape drug sensitivity patterns reveal vulnerable biological pathways. Heatmap showing the molecular pathways most sensitive to drug targeting for each screened sample from KEGG Pathways. Similar pathways are clustered together using the Jaccard distance and samples are clustered together by their Euclidian distance. Pathways are ranked independently for each sample based upon the results of drug screening experiments. Pathways targeted by the most effective drugs are ranked highest (red). Opaque squares indicate pathways in which more than 50% of the constituent genes were targeted in the drug panel. Pathways ranked in the top 50 for 20 or more samples are plotted. White squares indicate that the pathway was not targeted by any drugs in the screening experiments.

**Figure S6:**
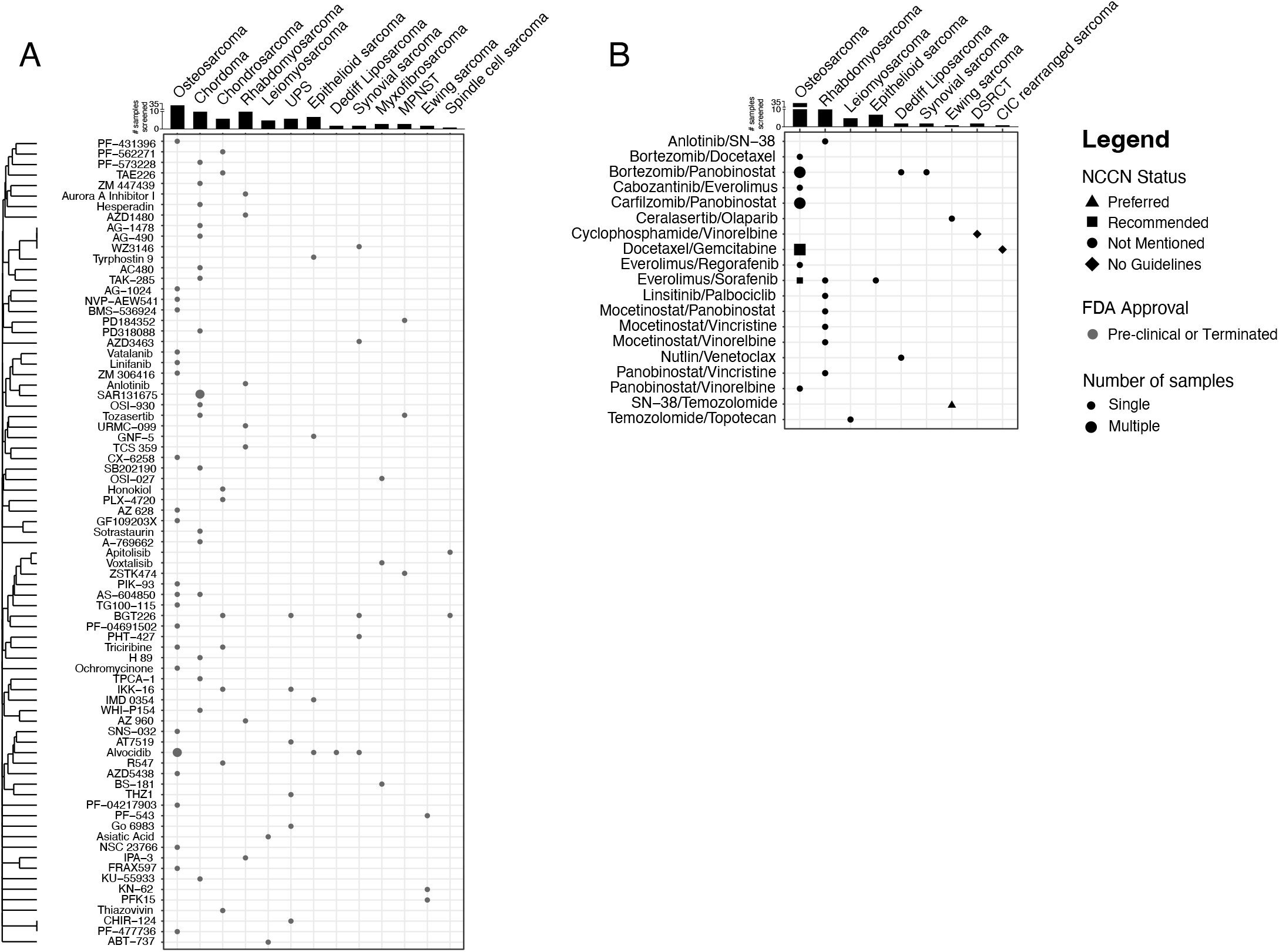
Drug availability and NCCN Guidelines status by histological subtype. We selected drug-diagnosis pairs of interest by cross-referencing the five most effective drugs for each sample with the 10% most responsive samples for each drug. (A) Single agent drugs that are not FDA approved are shown for each unique drug-diagnosis combination. The shape of the point indicates the current NCCN Guidelines for each drug. Triangles indicate drugs that are indicated as a preferred regimen. Squares signify drugs that are recommended for the subtype of interest and circles indicate that the drug is not currently discussed in the NCCN guidelines4,5. Diamond shape signifies that the histologic subtype has no guidelines, such as DSRCT and CIC rearranged sarcoma. The size of the marker indicates whether a single sample or multiple samples of a given histologic subtype was found to be among the five most effective for a drug. Drugs are clustered by similarity in gene targets using Jaccard distance. The number of samples screened for each histologic subtype is shown above. (B) Similar analysis is performed for combinational regimens and their NCCN recommendations across sarcoma histological subtype.

**Figure S7:**
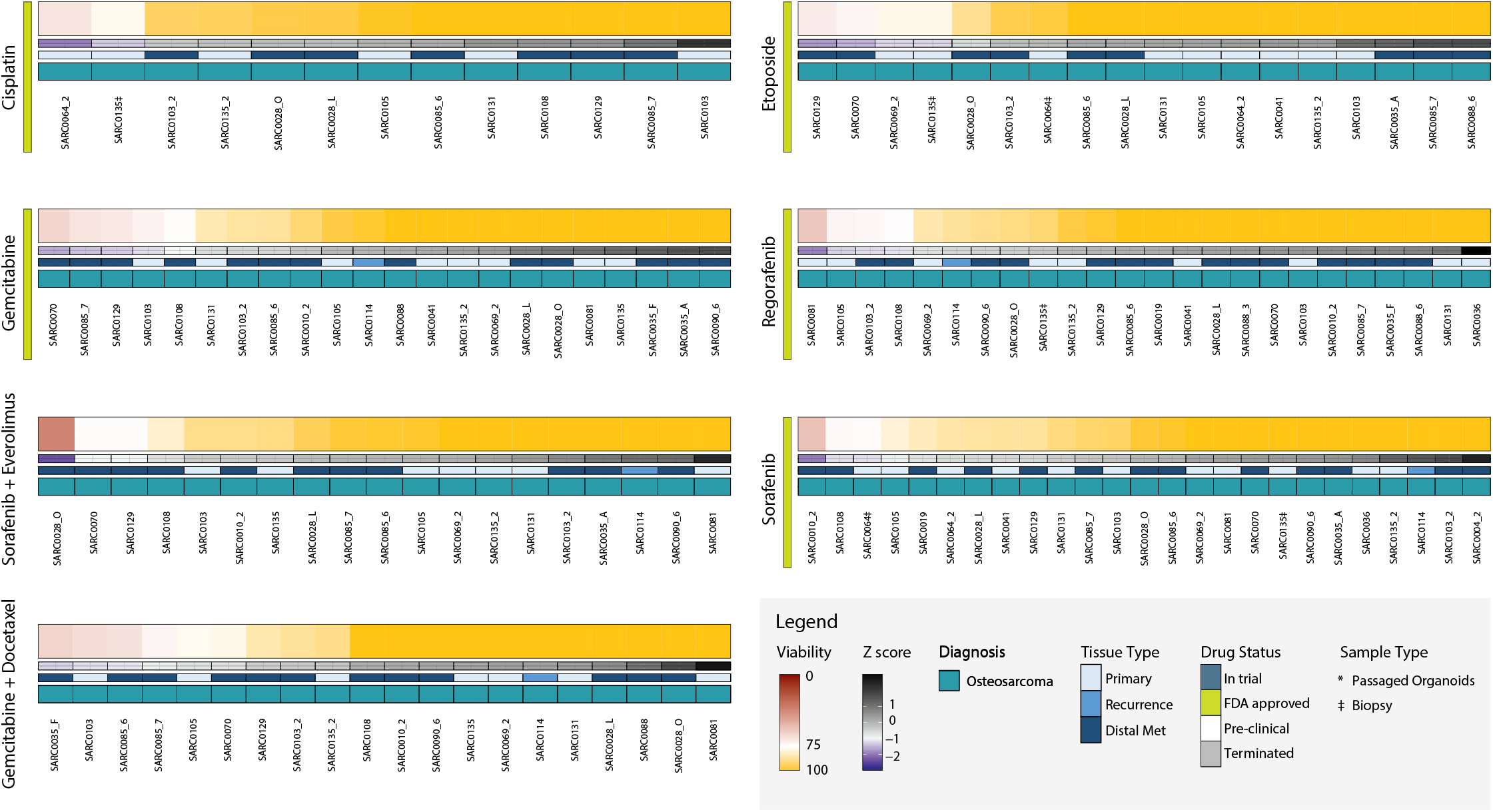
Osteosarcoma organoid sensitivity to treatment of NCCN recommended regimens. Heatmaps of organoid sensitivity to selected drugs from the NCCN recommendations were screened at 1 µM. Organoid viability for each sample is normalized to the mean organoid response to treatment with the selected drug across samples. Each column is a unique specimen, darker shades of red indicate greater sensitivity to treatment. Colored bars underneath each heatmap indicate the Z-score, lesion type, and diagnosis of each sample.

**Figure S8:**
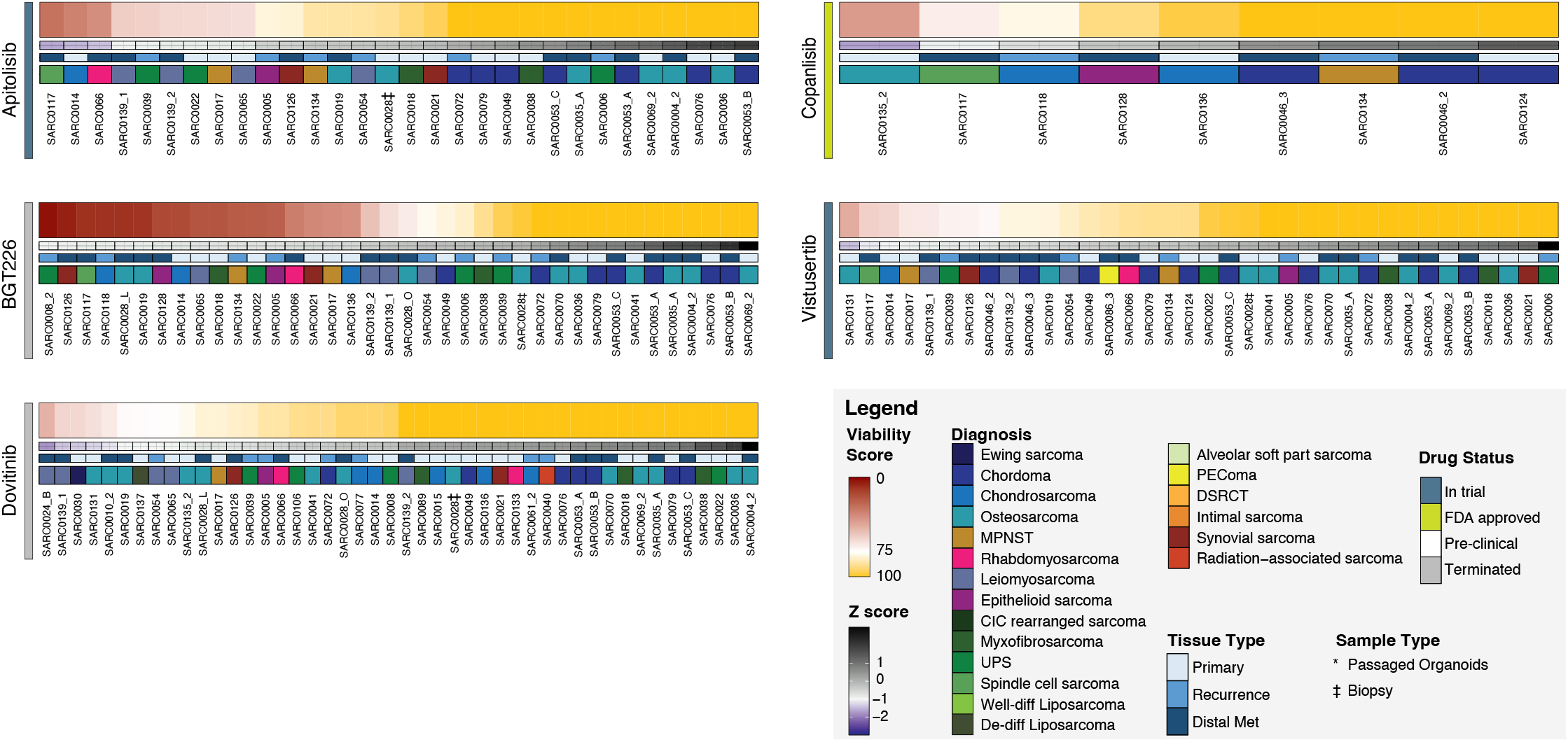
organoid sensitivity to treatment of mTOR/PI3K targeting drugs, dovitinib, apitolisib, copanlisib, BGT226, and vistusertib. Heatmaps of organoid sensitivity to selected drugs from the NCCN recommendations were screened at 1 µM. Organoid viability for each sample is normalized to the mean organoid response to treatment with the selected drug across samples. Each column is a unique specimen, darker shades of red indicate greater sensitivity to treatment. Colored bars underneath each heatmap indicate the Z-score, lesion type, and diagnosis of each sample.

